# The global evolutionary dynamics of natural capsule switching in a human-restricted bacterial pathogen

**DOI:** 10.64898/2026.09.24.752487

**Authors:** Chrispin Chaguza, Anna York, Stephanie W. Lo, Akuzike Kalizang’oma, Rebecca A. Gladstone, Brenda A. Kwambana-Adams, Rory Cave, Global Pneumococcal Sequencing (GPS) consortium, Keith P. Klugman, Cheryl P. Andam, Robert S. Heyderman, Lesley McGee, Jason W. Rosch, Nicholas J. Croucher, Stephen D. Bentley, Daniel M. Weinberger

**Author notes:** Correspondence (C.C.) and (D.M.W.).

## Abstract

Exchange of genetic loci encoding extracellular capsular polysaccharides in bacteria can change capsular antigens or serotype, which, in turn, can lead to evasion of vaccines targeting the capsule. However, the evolutionary dynamics and drivers of bacterial capsule switching remain poorly understood. Here, we investigate the evolutionary dynamics and correlates of capsule switching in bacteria, using the human-restricted bacterial pathogen *Streptococcus pneumoniae* as an example, using 10,558 genetically and geographically diverse whole-genome sequences. Our phylogenomic and ancestral state reconstruction analysis revealed that the propensity for capsule switching varies by capsule type and genetic background. We found capsule switches only between ∼4% of all possible capsule-type pairs, varying by specific pairs and switching direction. We also found associations between capsule switching patterns and genetic and ecological factors, including nasopharyngeal carriage duration, capsule biosynthesis locus size, and capsule thickness. Furthermore, the genetic similarity of the replacing and replaced capsule loci was primarily associated with the frequency of switching between specific capsule type pairs. Although a few specific non-capsule genomic loci were associated with the propensity for capsule switching across distinct lineages, the collective effect of the genetic background appears to influence capsule switching more strongly. These findings shed light on how bacterial capsule types and lineages undergo capsule switching influenced by genetic and ecological factors, informing our understanding of the emergence of novel capsule-genotype combinations that may impact the design of capsule-targeting glycoconjugate vaccines.

## Introduction

Extracellular capsule polysaccharide structures surround some bacterial cells (Cress et al. 2014; Whitfield et al. 2020). These capsules play a crucial role in physically protecting bacteria from desiccation and phage attack (Hamaguchi et al. 2018; Soundararajan et al. 2019), promoting virulence (Hathaway et al. 2016), and helping them evade the host immune system (Hyams et al. 2010; Liu et al. 2025). Genes that comprise capsule polysaccharide synthesis (*cps*) loci encode enzymes that synthesize these antigenically diverse capsules (Bentley et al. 2006; Mostowy and Holt 2018). These *cps* loci harbor an array of genes encoding proteins involved in sugar synthesis, assembly of lipid-linked polysaccharides, and export of mature sugars to the cell surface (Bentley et al. 2006; Mostowy and Holt 2018). These genetically diverse *cps* loci typically encode a single antigenically distinct capsule type (Bentley et al. 2006; Mostowy and Holt 2018). These capsules have become attractive targets for capsule-based vaccines, which induce capsule-specific immunity against targeted strains (Goldblatt 2000; Rappuoli et al. 2019). However, the evolutionary dynamics driving the emergence of novel capsule-genotype combinations that could evade current serotype-based vaccine formulations remain less understood.

Horizontal gene transfer (HGT) and recombination play a significant role in driving bacterial evolution, particularly in naturally transformable species (Chewapreecha et al. 2014; Arnold et al. 2022). The *cps* locus represents a major hotspot for genetic exchange in bacterial genomes (Mostowy et al. 2017). Such genetic exchanges (Mavroidi et al. 2007; Mostowy et al. 2014) alter the encoded capsules, leading to capsule switching (Moore et al. 2008). Consequently, the same genetic background or lineage can have variants that express different capsule types (Joshi et al. 2020). Furthermore, the same capsule types may be associated with multiple genetic backgrounds. Whole-genome sequencing (WGS) has revealed capsule switching in numerous pathogens, including *Streptococcus pneumoniae* (Brueggemann et al. 2007; Croucher et al. 2011; Wyres et al. 2013), *Streptococcus agalactiae* (Coffey et al. 1998; Bellais et al. 2012)*, Klebsiella pneumoniae* (Wyres et al. 2015), and *Streptococcus suis* (Murray et al. 2023). In bacterial species such as *S. pneumoniae* (pneumococcus), which kills more than 300,000 children globally (Wahl et al. 2018), capsule switching has led to variants that can evade vaccine-derived immunity targeting the capsule (Brueggemann et al. 2007). While capsule switching is a natural phenomenon, the widespread rollout of vaccines targeting specific capsule types, such as pneumococcal conjugate vaccines (PCVs) (21, 27), has raised fears about potential clonal expansion of capsule-switch mutants (Lucidarme et al. 2017).

Early studies of capsule switching primarily identified switches within a single genetic background (Alcalá et al. 2002; King et al. 2002; Tyler and Tsang 2004; Lancellotti et al. 2006; Brueggemann et al. 2007; Martins et al. 2010; Croucher et al. 2011; Bellais et al. 2012; Wyres et al. 2013; Chewapreecha et al. 2014; Croucher, Kagedan, et al. 2015; Wyres et al. 2015; Chaguza et al. 2017; Khan et al. 2022; Murray et al. 2023; Wang et al. 2023). These studies defined capsule switching as the expression of multiple capsule types within different isolates of the same genetic background, as determined by sequence type (ST) using multilocus sequence typing (MLST) (Jefferies et al. 2004; Croucher et al. 2011; Chewapreecha et al. 2014; Chaguza et al. 2017). Although this approach is simple, intuitive, and widely adopted globally because of MLST’s portability (Maiden et al. 2013), it has several limitations. First, MLST detects only recent capsule-switching events because its lower resolution compared with a whole-genome approach misses switches between similar strains with different or incorrect STs. Second, the MLST-based approach cannot infer the directionality of capsule switches. These limitations highlight the need for whole-genome sequencing (WGS) to investigate bacterial capsule-switching patterns with greater resolution (Chewapreecha et al. 2014; Croucher, Kagedan, et al. 2015; Joshi et al. 2020). Despite the increasing affordability of bacterial WGS (Blackwell et al. 2021), most studies have examined capsule switching in individual lineages, used small datasets, or sampled isolates from one or a few countries (Jefferies et al. 2004; Brueggemann et al. 2007; Croucher et al. 2011; Golubchik et al. 2012; Croucher et al. 2013; Croucher, Kagedan, et al. 2015; Chaguza et al. 2017; Khan et al. 2022). Because of the remarkable genetic and geographical diversity of particular bacterial species (13, 41, 42), sequencing extensive, globally representative strain collections is necessary to gain robust evolutionary insights into capsule switching. Such sequencing studies may pinpoint capsule types and genetic backgrounds more prone to capsule switching, which can drive the emergence of vaccine-escape strains when a capsule type couples with an optimal genetic background (Brueggemann et al. 2007; Golubchik et al. 2012).

In this study, we investigate the evolutionary dynamics and correlates of capsule switching in a global bacterial population. We use the clinically significant human-restricted commensal and pathogenic bacterium *S. pneumoniae*, also known as pneumococcus, as an example. We performed a computationally intensive, large-scale phylogenomic analysis of 10,558 whole-genome sequenced nasopharyngeal and disease pneumococcal isolates, carefully selected from our globally sampled genome collection generated by the Global Pneumococcal Sequencing (GPS) consortium (43), to ensure representation by capsule type, genetic background, and country of origin. We first quantified and assessed the variability in the propensity of capsule types and genetic backgrounds or lineages to undergo capsule switching, as well as the frequency with which specific capsule types were acquired and lost within the global phylogeny. We then estimated the proportion and frequency of direct capsule switching between all possible combinations of capsule types expressed in the global pneumococcal phylogeny. Lastly, we assessed factors associated with capsule-switching patterns across different bacterial genetic backgrounds and between specific capsule-type pairs. Understanding these evolutionary dynamics could provide critical insights with implications for predicting long-term capsule-type dynamics and designing capsule-based vaccines to prevent infections with novel capsule types (Mostowy and Holt 2018).

## Results

### Capsule switching varies by specific pneumococcal capsule type and genetic background

The pneumococcal *cps* locus is a known evolutionary hotspot for HGT and recombination (Mostowy et al. 2017), and switching at this locus can lead to the expression of a single capsule type across multiple genetic backgrounds or lineages (Mostowy et al. 2017). We hypothesized that capsule-switching propensity varies by capsule type and genetic background, reflecting the influence of both capsule and lineage on bacterial capsule switching in the pneumococcal population. To assess this, we compared capsule-type distributions across pneumococcal lineages, defined using the global pneumococcal sequence cluster (GPSC) nomenclature (43), to quantify the frequency of capsule-type switching between genetic backgrounds. To ensure robust detection of potential capsule switching events, we leveraged a large and geographically diverse global collection of 10,558 pneumococcal genomes (from 56 countries and six continents) carefully sub-sampled from a larger dataset of 21,155 genomes generated by the Global Pneumococcal Sequencing (GPS) project (Gladstone et al. 2019) **(Fig. 1A-C and Supplementary Data 1)**. Our dataset included genomes associated with each capsule type, genetic background, and country, including those sampled at low frequency in a few localized settings to capture potential rare capsule-switching events, as well as those present in oversampled, highly prevalent lineages (Gladstone et al. 2019). These genomes enabled an extensive investigation of capsule-switching patterns in the global pneumococcal population.

**Fig. 1.**
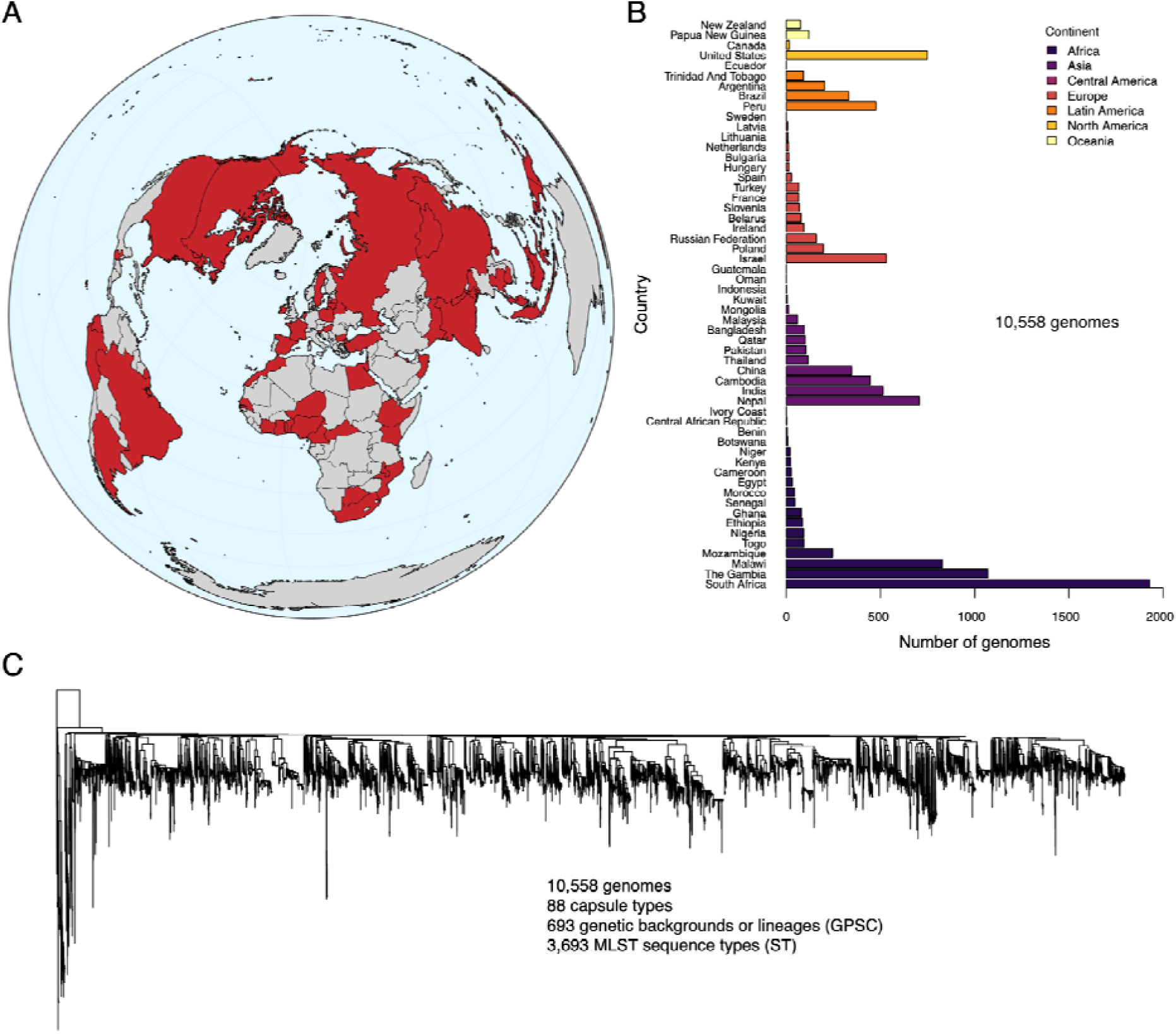
A geographically and genetically diverse dataset of whole-genome-sequenced S. pneumoniae strains facilitates global population-level analysis of capsule-switching patterns. **(A)** A world map showing the origin (red color) of the pneumococcal isolates sequenced in this study through the Global Pneumococcal Sequencing (GPS) project (https://www.pneumogen.net/gps/). This study used sequenced genomes from 56 countries across seven continents. In total, 88 capsule types (serotypes), 3,693 known sequence types (ST) based on the pneumococcal multilocus sequence typing (MLST) scheme, and 693 known genetic backgrounds or lineages defined using the Global Pneumococcal Sequence Cluster (GPSC) nomenclature. The full dataset of pneumococcal genomes and metadata used in this study is available from the GPS Monocle website (https://data.monocle.sanger.ac.uk/). **(B)** Total number of pneumococcal genomes selected from each country. The genomes included in this study were selected from a larger GPS dataset to include up to five genomes for each combination of genetic backgrounds defined by the GPSC and MLST, capsule type, and country to ensure the inclusion of every pneumococcal strain from all the sampled countries, including those found at low frequencies (see methods). **(C)** A global whole-genome maximum-likelihood pneumococcal phylogeny showing genetic relatedness of the genomes included in this study. We generated the phylogeny from whole-genome multiple sequence alignments of consensus sequences created by mapping short sequence reads from each isolate to a single reference sequence (see Methods). We included a single Streptococcus mitis genome as an outgroup to root the phylogeny. The full list of genomes used in this study is available in **Supplementary Data 1**.

We first assessed how often pneumococcal capsule types were acquired across different genetic backgrounds. We observed an unequal distribution of capsule types across different pneumococcal lineages, reflecting varying acquisition patterns within distinct genetic backgrounds **(Fig. 2A, B, Supplementary Fig. 2, 3, and Supplementary Data 2)**. Most lineages expressed multiple capsule types, indicating a rich history of capsule switching depending on capsule type. The capsule types expressed on the most lineages were 6A, 19A, 6B, 19F, and 23F **(Fig. 2A, B, Supplementary Fig. 2, 3, and Supplementary Data 2)**. In contrast, capsule types, such as 1, 2, and 11E, were expressed in only a single lineage each **(Supplementary Fig. 2C and Supplementary Data 2)**.

**Fig. 2.**
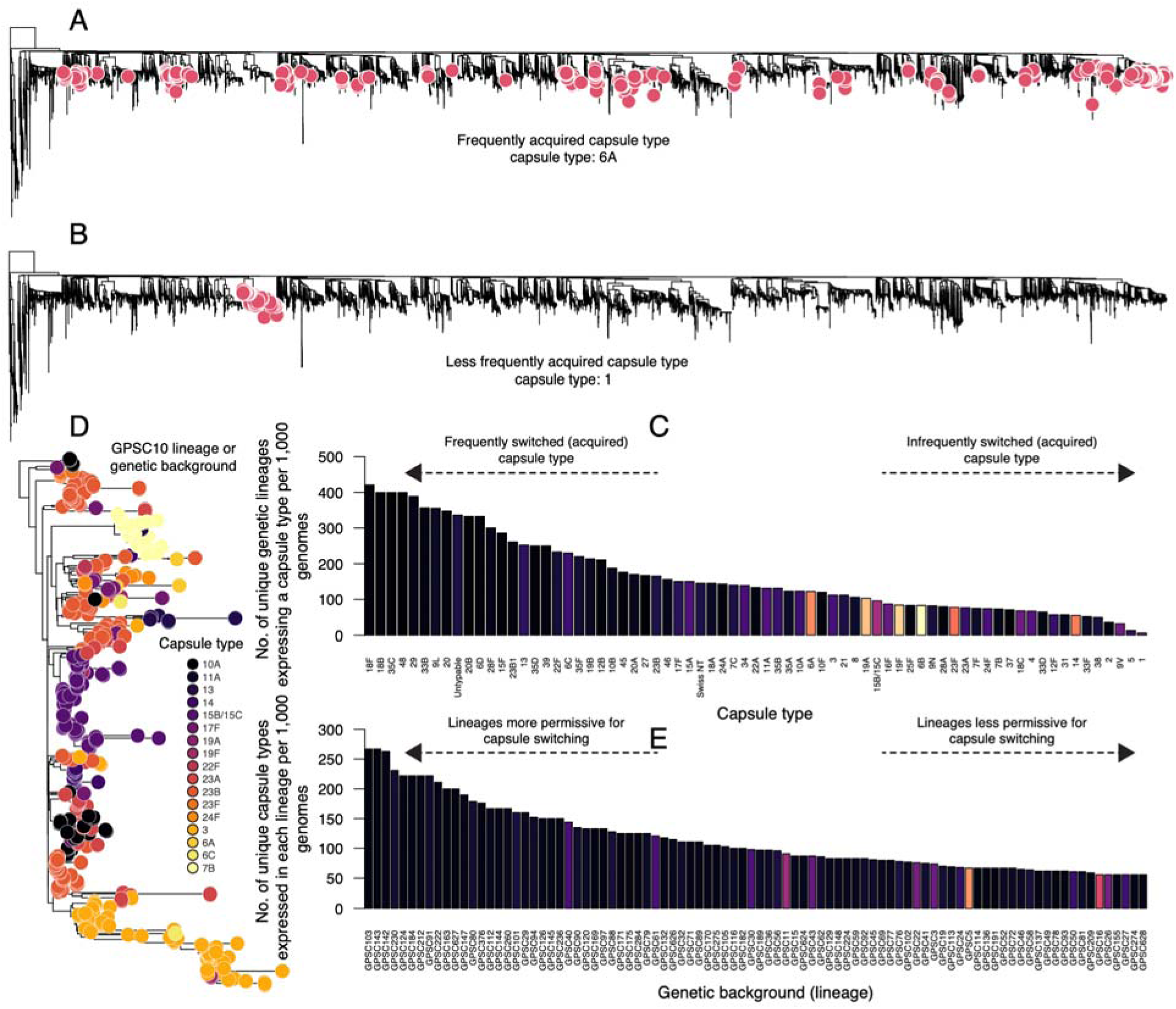
The distribution of capsule types across lineages in the global pneumococcal phylogeny provides insights into the propensity for capsule switching by capsule type and genetic background or lineage. **(A)** A global whole-genome maximum-likelihood phylogeny illustrates the distribution of pneumococcal strains expressing the type 6A capsule across genetic backgrounds or lineages. The widespread presence of 6A strains across the phylogeny indicates that this capsule is readily acquired through capsule switching. **(B)** A maximum-likelihood phylogeny showing the distribution of capsule type 1 strains across diverse genetic backgrounds (i.e., GPSC lineages). The single lineage expressing type 1 capsule suggests infrequent capsule switching involving this capsule and lineage in the phylogeny. The phylogenies in panels A) and B) were generated from the whole-genome multiple sequence alignments of consensus sequences generated by mapping short sequence reads of each isolate to a single reference sequence (see methods). We included a single genome from Streptococcus mitis (not shown in the phylogeny) as an outgroup to root the phylogeny. **(C)** The bar chart shows the adjusted number of distinct genetic backgrounds that express a particular capsule type per 1,000 genomes. Supplementary Data 2 provides the specific counts shown in the bar chart. **(D)** A maximum likelihood phylogeny showing an example of a GPSC lineage expressing many capsule types indicates a genetic background potentially more permissive to capsule switching. **(E)** Bar chart showing the adjusted number of unique capsule types expressed in a single genetic background or lineage per 1,000 genomes. For clarity, the bar chart shows only the top 50 genetic backgrounds or GPSCs expressing the most capsule types. The complete list of genetic backgrounds and the number of unique capsule types they contain is available in **Supplementary Data 4**. Panels C and E are based on only capsule types and lineages with at least ten sequenced genomes in the dataset, and the color of the bars in the bar plots corresponds to the number of genomes included in the analysis for the capsule type or lineage – the lighter the color, the more genomes **(Supplementary Data 1)**.

To account for sampling bias due to the presence of more commonly sampled capsule types in more lineages than rarely sampled types, we adjusted the estimates for the number of lineages expressing each capsule type (capsule acquisition frequency) to reflect the number of genomes included in the analysis for each capsule type **(Supplementary Fig. 2A and 2B)**. Based on the adjusted estimates, we observed that some capsule types are more readily acquired than others across different lineages. Capsule types found across the most lineages included 18F, 18B, 35C, 29, and 33B **(Fig. 2B, Supplementary Data 3, and Supplementary Fig. 4A)**. Interestingly, despite the importance of the capsule for pneumococcal colonization and virulence (Kadioglu et al. 2008), untypable strains harboring a partial or complete deletion of the *cps* loci were among the top ten capsule-types expressed across different genetic backgrounds, suggesting that loss of encapsulation may be transiently evolutionarily advantageous to the pneumococcus (Kadioglu et al. 2008; Andam and Hanage 2015). We then assessed the propensity of different pneumococcal genetic backgrounds to undergo capsule switching. We reasoned that genetic backgrounds more permissive to capsule switching would harbor more capsule types than more restrictive ones. After accounting for the number of genomes sampled per lineage, we observed that the frequency of expressed capsule types varies across different genetic backgrounds **(Fig. 2C, D; Supplementary Fig. 4B-D; and Supplementary Data 4 and 5)**.

Altogether, these findings show that the propensity for capsule switching varies not only across capsule types but also across genetic backgrounds, hinting at potential genetic and ecological barriers to natural capsule switching in evolving pneumococcal populations.

### Ancestral reconstruction reveals patterns of pneumococcal capsule switching

We then quantified capsule acquisition and loss across the global phylogeny using ancestral state reconstruction for each capsule type. This approach captures independent acquisitions and losses of capsule types within a single lineage by directly considering the evolutionary history of the strains **(Fig. 3A and Supplementary Fig. 6)**. Interestingly, the capsule acquisition frequency using the ancestral state reconstruction approach revealed a strong correlation with the crude adjusted estimates of capsule acquisition based on counting the number of unique lineages expressing each capsule type – the non-ancestral state reconstruction approach (Pearson correlation coefficient=0.96, *P*<2.2×10^-16^) **(Supplementary Fig. 5A, B)**. However, the non-ancestral state reconstruction approach slightly underestimated the frequency of capsule acquisition compared with the ancestral state reconstruction method, likely due to convergent evolution or homoplasy (Wilcoxon signed-rank test, *P*=5.83×10^-05^) **(Supplementary Data 6, 7)**. Therefore, ancestral state reconstruction improved detection of capsule acquisitions by accurately capturing instances where the same capsule type is acquired multiple times within the same genetic background.

**Fig. 3.**
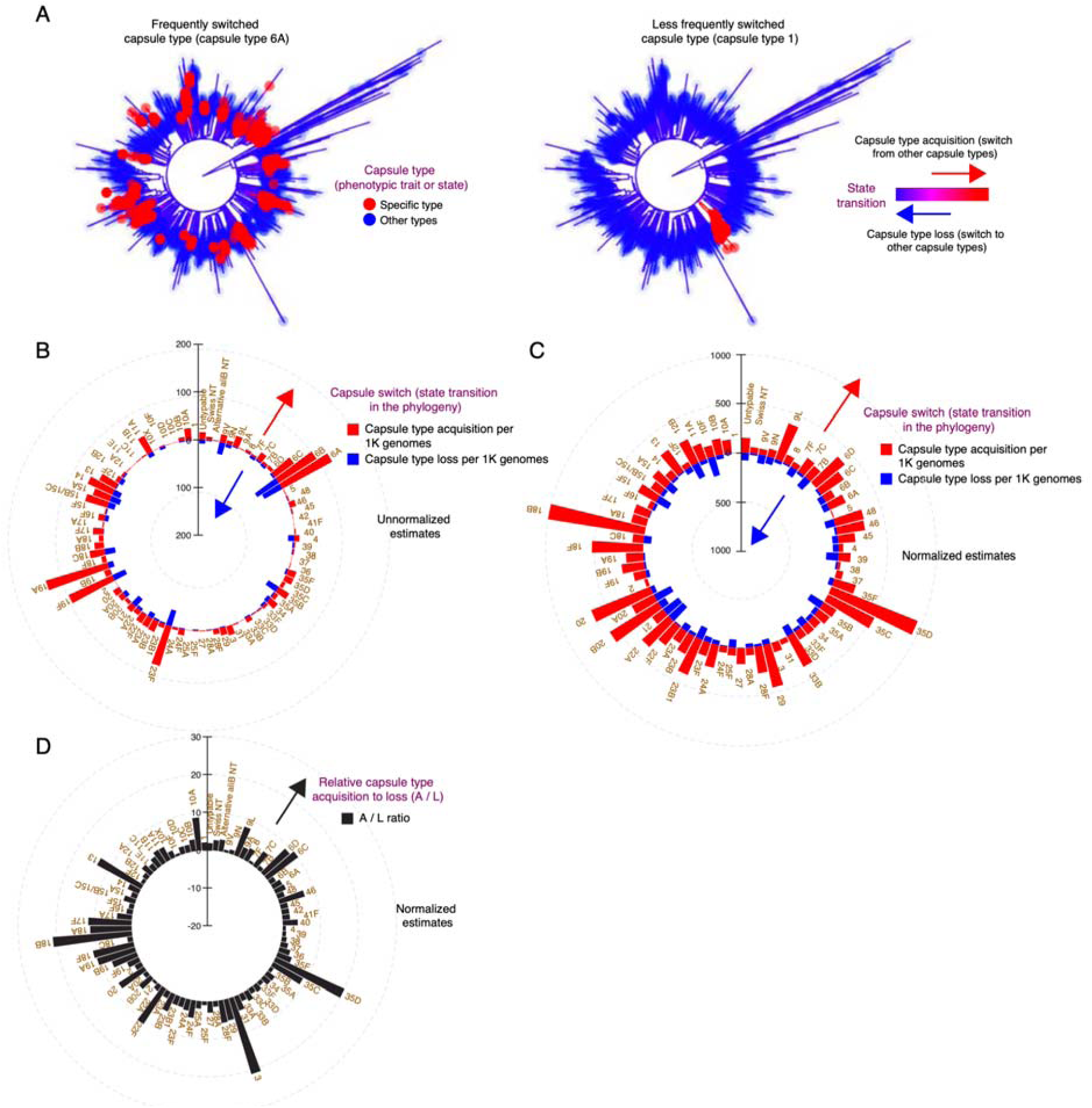
Mapping the evolution of individual capsule types across the whole-genome-based pneumococcal phylogeny using ancestral state reconstruction robustly quantifies the frequency of capsule switching patterns globally. **(A)** A rooted global pneumococcal whole-genome maximum-likelihood phylogeny showing reconstructed capsule acquisition and loss patterns. A capsule acquisition event corresponds to a capsule switch from other capsule types to a particular type, while a capsule replacement event represents a switch from a particular capsule type to others. Two capsule types are shown: one associated with longer carriage duration and high prevalence in nasopharyngeal carriage, frequently acquired across the global phylogeny (capsule type 6A), and one associated with short carriage duration and low prevalence in nasopharyngeal carriage (capsule type 1). Phylogenies for the other capsule types are shown in Supplementary Fig. 6. **(B)** A polar histogram summarizes the acquisition and loss frequency of all capsule types across the global pneumococcal phylogeny. **(C)** A polar histogram summarizing the acquisition-to-loss ratio of all capsule types across the global pneumococcal phylogeny. We calculated the capsule acquisition-to-loss ratio for all capsule types by adding pseudocounts of 1 to both the numerator and denominator to avoid division by zero when no capsule replacements were observed. **(D)** A polar histogram summarizing the frequency of acquisition and loss of all capsule types across the global pneumococcal phylogeny, based on estimates from the number of genomes for each capsule type included in the dataset. We express adjusted capsule-type acquisition and loss frequency as the number of capsule-type acquisitions or losses per 1,000 genomes, calculated only for capsule types with at least 10 genomes in the dataset. We calculated the capsule acquisition-to-loss ratio for all capsule types by adding pseudocounts of 1 to both the numerator and denominator to avoid division by zero when no capsule replacements were observed.

We then applied ancestral state reconstruction to assess the acquisition and loss patterns of different capsule types in the global phylogeny. We defined capsule acquisition as switching to a specific capsule type, while capsule replacement reflected switching from one capsule type to another. Similarly, we observed that different capsule types were acquired and lost at unequal frequencies in the pneumococcal phylogeny **(Fig. 3B, C, and Supplementary Fig. 6)**. We then compared whether each capsule type was acquired at the same frequency as it was lost across the phylogeny. We found that most capsule types were acquired more often than they were lost (Wilcoxon signed-rank test, *P=*1.09×10^-11^). We then compared the ratio of capsule types acquired to those lost across the phylogeny. We found that at a higher acquisitions-to-loss ratio (A/L ratio) for capsule type 3 (A/L ratio: 21.0), 18B (A/L ratio: 21.0), 35D (A/L ratio: 20.0), and 13 (A/L ratio: 13.33), which indicated that these capsule types tended to replace other capsule types than other types replaced them after they were acquired across the global pneumococcal phylogeny **(Fig. 3D)**. Further analysis of capsule type 3, which expresses the thickest pneumococcal capsule(Paton and Trappetti 2019), revealed 20 independent acquisitions across the global pneumococcal phylogeny, with no losses **(Fig. 3B, C, and Supplementary Data 6, 7)**. Likewise, type 1 capsule, which is rarely carried in the nasopharynx (Ritchie et al. 2012), an ecological niche where interactions and genetic exchange occur between strains (Marks et al. 2012), was acquired once in the phylogeny and showed no switches to other capsule types. We speculated that the lack of switching among type 1 strains reflected a reduced permissiveness of their genetic background to capsule switching, since transformation varies by more than 100-fold among strains (Evans and Rozen 2013), which in this case may be partly driven by a prophage that inhibits transformation (Kwun et al. 2022).

Collectively, these findings demonstrate that the acquisition and loss of pneumococcal capsule types in the global phylogeny occur at unequal frequencies, potentially indicating the influence of unknown capsule-dependent factors on capsule switching.

### Only a minority of capsule-type pairs show evidence of capsule-switching

Previous studies have primarily focused on capsule switching within specific clones (Jefferies et al. 2004; Pillai et al. 2009; Croucher, Kagedan, et al. 2015; Gladstone et al. 2015; Golden et al. 2015; Sheppard et al. 2019; Chaguza, Heinsbroek, et al. 2020), limiting our understanding of these patterns across the entire pneumococcal species. Consequently, which capsule type pairs show evidence of detectable switching, and to what extent, remains unknown. We hypothesized that only a small fraction of capsule-type pairs show evidence of direct recent or historical capsule switching because genetic and ecological constraints limit recombination. We performed ancestral state reconstruction of each capsule-type pair as discrete ancestral states in the phylogeny to infer the number and directionality of observable switches between capsule types. We analyzed 7,656 possible capsule-type pairs based on 88 capsule types in our global dataset to infer the frequency and directionality of reconstructed capsule or state transitions in the phylogeny for each pair. We found that 4.36% (334 of 7,536) of possible capsule switch pairs had a representative switch evident in our dataset, suggesting that a minority of pneumococcal capsule-type pairs had a switch that successfully propagated in the population **(Fig. 4A-C and Supplementary Data 8, 9)**. Overall, these findings suggest that only a minority of pneumococcal capsule-type pairs have undergone capsule switching, further highlighting potential barriers to capsule switching and potentially lower fitness and survival of capsule-switch mutants compared to the wild-type parental strains within the pneumococcal population.

**Fig. 4.**
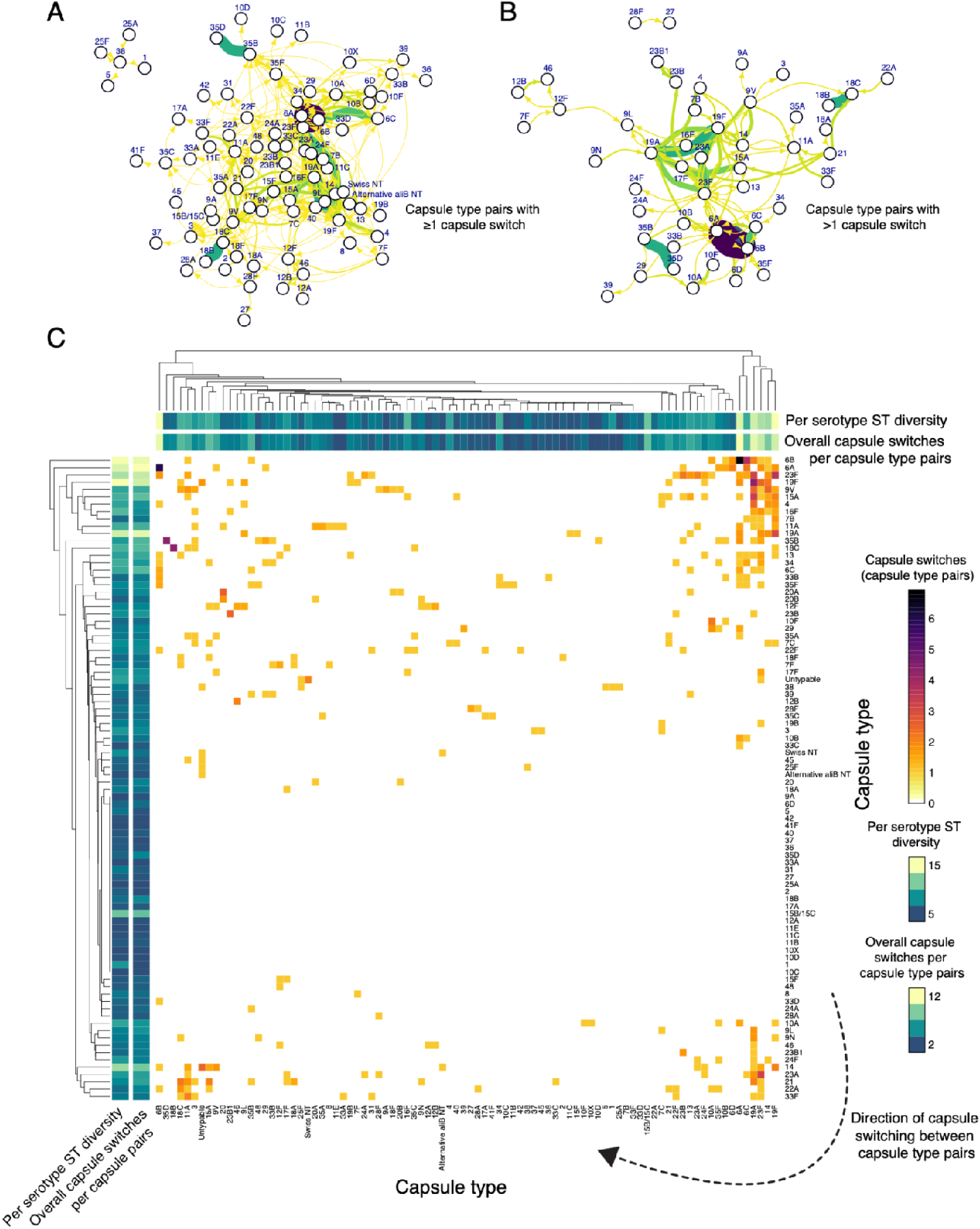
Capsule switching between all possible capsule-type pairs in the global pneumococcal strain collection. **(A)** A directed network showing capsule switching events between pairs of capsule types inferred using the ancestral state reconstruction approach. Nodes represent capsule types, and edges indicate capsule-switch events observed between pairs of capsule types. Edge thickness is proportional to the observed frequency of capsule-switching events, without adjusting for the number of genomes associated with each capsule type in the analyzed dataset. The arrows on each network edge show the inferred direction of capsule-switching events. Edge colors are assigned arbitrarily to visualize overlapping edges. **(B)** A subnetwork showing capsule switches identified only between capsule pairs with at least two capsule-switch events. **(C)** A heatmap showing the inferred number of capsule-switching events between capsule-type pairs. The color intensity in the heatmap corresponds to the inferred number of transitions or switches from one capsule type to another. The horizontal and vertical color strips located at the tips of the dendrogram at the top and left side of the heatmap show the “population-level capsule type diversity”, reflecting the number of unique MLST sequence types (STs) associated with a single capsule type as a proxy for the distribution or prevalence of the capsule type. Essentially, capsule types that are highly prevalent and have circulated in the population for extended periods are more likely to have had more opportunities to diversify genetically, leading to more STs and, therefore, a higher population-level diversity value than recently emerged and less prevalent capsule types globally. The overall number of capsule switches corresponds to the total observed number of capsule switches, either switching from other types to the capsule type (gain) or from the capsule type to another type (loss), associated with each capsule type (see Fig. 3), besides those inferred between specific pairs of capsule types. The frequency of capsule switching shown in the figure key is square root transformed for visual clarity.

### The frequency of capsule switching varies depending on the specific capsule type pairs

Next, we investigated the frequency and directionality of capsule switching between specific capsule-type pairs. Because of genetic and ecological barriers to recombination (30, 56, 57), we anticipated that switching frequency and directionality would vary across pneumococcal capsule-type pairs. Based on the ancestral state reconstruction analysis, we found the highest frequency of capsule switches between commonly carried capsule types (Chaguza, Senghore, et al. 2020), including 6A→6B (n=83), 19A→19F (n=28), 18B→18C (n=19), 19F→23F (n=19), 35B→35D (n=19), 6B→6C (n=18), and 19A→23F (n=14) **(Fig. 4A-C and Supplementary Data 8,9)**. We observed more frequent capsule switching within serogroups than between serogroups. We then assessed whether capsule-switching frequency between capsule-type pairs was symmetrical in each direction. Interestingly, within this dataset, we observed asymmetry in capsule-switching frequency between specific capsule-type pairs, with some switches occurring more often in one direction than the other **(Fig. 4A-C and Supplementary Data 8, 9)**. Examples of capsule type pairs showing pronounced asymmetry in the frequency of capsule switches included 35B and 35D (35B→35D: n=19; 35D→35B: n=0), 6B and 6C (6B→6C: 14; 6B→6C: 2), and 15A and 19A (15A→19A: 11; 19A→15A: 0), 18B and 18C (18B→18C: 0; 18C→18B: 19), 6A and 6B (6A→6B: 36; 6B→6A: 48), and 19A and 19F (19A→19F: 9; 19F→19A: 19). Overall, the variability and asymmetry in the capsule-switching frequency between pneumococcal capsule-type pairs further highlighted the potential barriers to capsule switching in pneumococcus.

### Capsule type acquisition and loss in the global pneumococcal phylogeny correlate with nasopharyngeal carriage properties

Having observed unequal frequencies of capsule acquisition, loss, and direct switches between specific capsule pairs, we next assessed how specific genetic and ecological factors influence pneumococcal capsule-switching patterns. We hypothesized that capsule-associated factors, including nasopharyngeal carriage prevalence (Chaguza et al. 2016), cell-to-cell interaction (Cowley et al. 2018), competition (Trzciński et al. 2015), extracellular capsule thickness (Yother et al. 1986; Weiser and Kapoor 1999; Pearce et al. 2002; Chewapreecha et al. 2014), and *cps* loci similarity (Mavroidi et al. 2007; Croucher, Kagedan, et al. 2015), may influence capsule switching patterns. Because we observed a positive association between acquisition frequency and the number of genomes associated with each capsule type, we adjusted capsule-switching estimates, as oversampled capsule types may show more switching than undersampled ones **(Supplementary Fig. 1B)**.

Univariate generalized linear regression analysis based on these adjusted estimates revealed a positive association between capsule acquisition frequency and carriage duration (β=0.16, *P<*2×10^-16^) **(Fig. 5A and Supplementary Data 10)**. Because previous studies suggest that shorter DNA fragments recombine more frequently than longer fragments (19, 64), we postulated that cps locus length may influence capsule acquisition or switching frequency. Repeating the regression analysis to include the *cps* locus length as an additional covariate revealed a positive association of the capsule acquisition frequency with carriage duration (β=0.48, *P<*2×10^-16^) and a negative association with *cps* locus length (β=-0.21, *P<*8.79×10^-13^) **(Fig. 5A, Supplementary Fig. 7, and Supplementary Data 10)**. The results were consistent with the notion that shorter *cps* loci are more readily acquired than longer *cps* loci. Since the capsule inhibits pneumococcal DNA uptake and transformation (13, 61–63), we further analyzed a smaller subset of capsule types (n*=*15) with at least ten available genomes and known capsule thickness measurements (Weinberger et al. 2009). Again, this analysis showed a similar association between capsule acquisition frequency and carriage duration (β=0.21, *P<*3.54×10^-09^) and *cps* locus length (β=-0.36, *P<*2.84×10^-13^) but a positive association with capsule thickness (β=0.40, *P<*2×10^-16^) **(Fig. 5A and Supplementary Data 10)**.

**Fig. 5.**
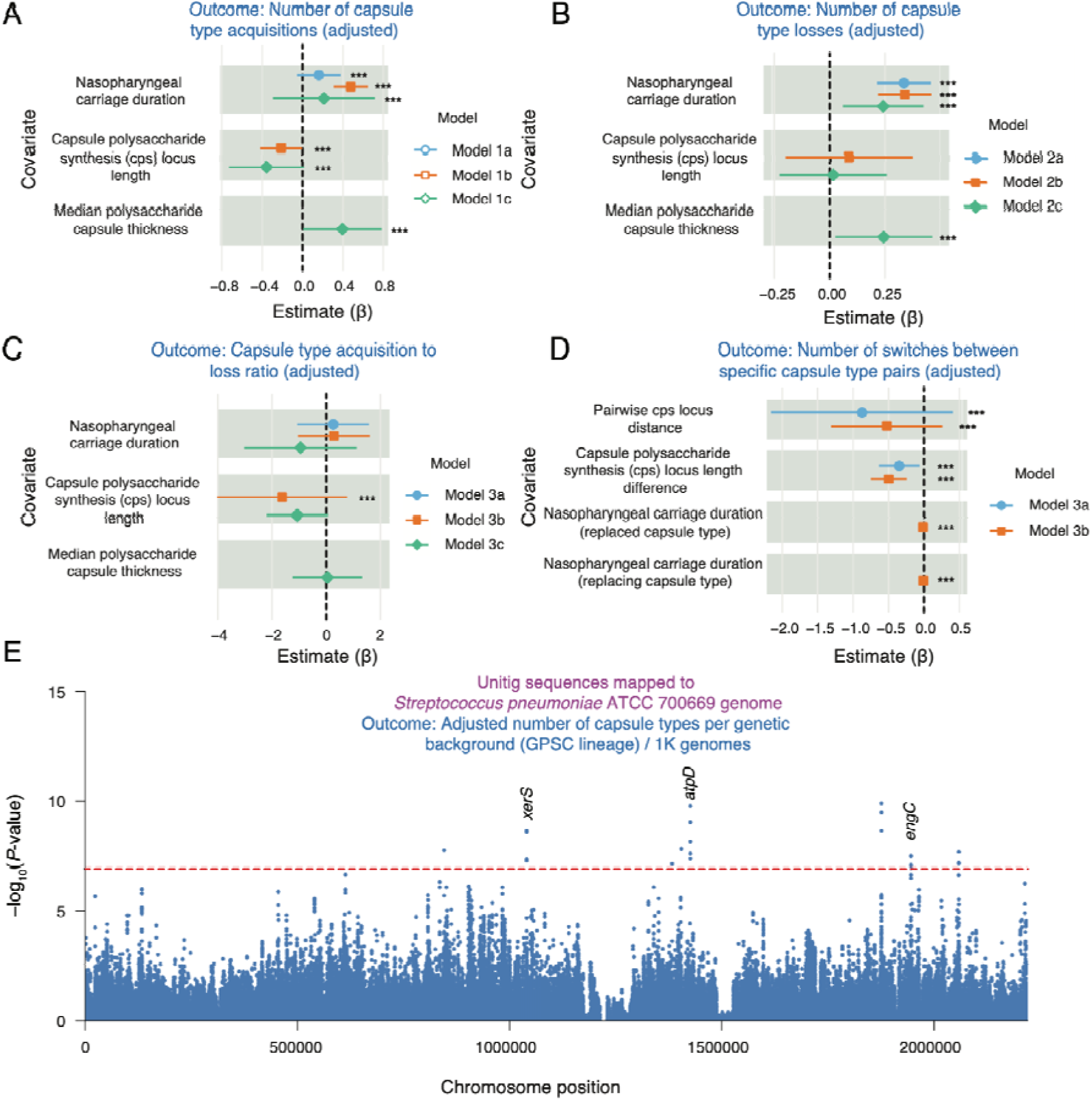
Genetic and non-genetic factors are associated with capsule switching in the global phylogeny. **(A)** Summary of the Poisson generalized linear regression model coefficients showing the association between genetic and non-genetic factors and the number of acquisitions of each capsule type (switches from other capsule types to a specific capsule type). **(B)** Summary of Poisson generalized linear regression model showing the association between genetic and non-genetic factors with the number of losses of each capsule type (switches from a specific capsule type to other capsule types). **(C)** Summary of Poisson generalized linear regression model showing the association between genetic and non-genetic factors with the ratio of the frequency of acquiring to losing each capsule type (A/L ratio). **(D)** Summary of Poisson generalized linear regression model showing the association between genetic and non-genetic factors with the number of capsule switches between specific pairs of capsule types. The asterisks in panels A-D show covariates with a statistically significant association with the response variable. **(E)** Manhattan plot showing the statistical significance from a GWAS analysis using Pyseer(Lees et al. 2018) of the association between genomic loci (unitigs) and the propensity of each genetic background or GPSC lineage to capsule switching, defined as the number of unique capsule types expressed in each lineage per 1,000 genomes **(see** Fig. 2E). The statistical significance (*P*-value) was calculated using the likelihood ratio test. To annotate the variable-length *k-mer* sequences or unitigs, each sequence was mapped against the ATCC 700669 pneumococcal reference genome (GenBank accession: NC_011900 (Croucher et al. 2009)) to visualize their genomic context in the pneumococcal genome and identify genomic features, including genes, tRNA, rRNA, etc., present in the regions containing the sequences. Each unitig location is represented by a single blue dot in the Manhattan plots. The horizontal red dotted line in the Manhattan plot marks the genome-wide threshold for considering a unitig sequence significantly associated with the propensity for capsule switching in lineages. Unitigs with statistical significance above the red line were deemed statistically associated with the genetic background’s propensity to capsule switching. Known gene names harboring unitig sequences with statistical significance values above the genome-wide threshold are labeled next to the unitig sequences.

Next, we performed similar regression analyses to assess associations between carriage duration, cps locus size, capsule thickness, and capsule replacement frequency. A regression model with these three covariates similarly showed a positive association between capsule replacement frequency and carriage duration (β=0.24, *P<*1.45×10^-05^) and a negative association with capsule thickness (β=0.24, *P=*0.0002) but no association with the *cps* locus size (β=0.12, *P*=0.811) **(Fig. 5B and Supplementary Data 10)**. In addition, we assessed the association between these three covariates and the capsule-acquisition-to-loss ratio (A/L ratio) to determine factors that may influence the relative permissiveness of capsule types to replace or be replaced by other capsule types. We generally found no statistically significant association between these covariates and the capsule A/L ratio, except for one model, which showed a negative association between the *cps* locus length and the A/L ratio **(Fig. 5C and Supplementary Data 10)**. Finally, we assessed the association between carriage duration, the difference in cps locus length, and cps locus similarity between replacing and replaced capsule types for pairs with available carriage duration data and at least ten genomes. Based on the saturated model, we found a negative association between the *cps* locus distance or dissimilarity (β=-0.52, *P<*0.0003) and *cps* length difference (β=-0.49, *P<*1.90×10^-09^) with the adjusted frequency of capsule switching between pairs of capsule types **(Fig. 5D and Supplementary Data 10)**, consistent with the notion that recombination decreases with increasing sequence dissimilarity (Majewski et al. 2000). Although we found an association between donor carriage prevalence (β=-0.01, P<2.08×10-10) and recipient capsule types (β=-0.01, P<2.70×10-05), the nearly negligible effect sizes suggest a smaller effect of carriage prevalence on the occurrence of direct capsule switches between capsule pairs.

These findings demonstrate a correlation between factors associated with the duration of nasopharyngeal carriage of pneumococcal capsule types and capsule-switching patterns in the pneumococcal population.

### The genetic background exerts a greater collective influence on capsule switching propensity than specific individual loci

Beyond capsule-related factors, variation in the pneumococcal genome, including genes that epistatically interact with the cps locus, may influence variability in capsule-switching patterns across lineages (Croucher, Kagedan, et al. 2015; Joshi et al. 2020). Because transformation rates vary across pneumococcal strains (49, 66) and specific loci appear to influence transformation in other bacterial pathogens (67, 68), we hypothesized that different pneumococcal lineages harbor specific genetic variation in competence machinery genes and other loci that hinder or promote the emergence of capsule-switched strains. We tested this hypothesis by performing a bacterial genome-wide association study (GWAS) using linear mixed-effects models to assess the association of specific genomic loci with each lineage’s propensity to undergo capsule switching, i.e., the adjusted number of unique capsule types expressed in each lineage **(Fig. 2D, E)**. Due to the high recombinogenic nature and genetic diversity of the pneumococcus (Croucher et al. 2011; Chewapreecha et al. 2014), our GWAS analysis compared the distribution of unitigs or variable-length *k-*mer sequences between representative strains from lineage to capture the effect of genomic variants within both the conserved (core) and non-conserved (accessory) genomic regions (Chewapreecha et al. 2014).

We found some evidence of statistical association between thirty-six unitigs and the propensity of different pneumococcal genetic backgrounds to capsule switching. These unitigs mapped to several genic and intergenic regions. However, contrary to our expectation, no unitigs mapped to known competence machinery genes. Instead, we found matches to the unitigs in genes associated with diverse functions, namely the F0F1 ATP synthase subunit gene (*atpD*; locus tag: SPN23F_14720), tRNA (locus tag: SPN23F_08650), rhomboid family membrane protein-encoding gene (locus tag: SPN23F_21180), ribosome-associated GTPase (*engC*; locus tag: SPN23F_20040), site-specific tyrosine recombinase (*xerS*; locus tag: SPN23F_10630), RNA helicase (locus tag: SPN23F_14460), SpoU rRNA methylase family gene (locus tag: SPN23F_14210) and some intergenic regions (**Fig. 5D and Supplementary Data 10)**. Of these genes, we observed statistical association in an annotated region (β=0.929, *P*=1.28×10^-10^), *atpD* (β=-0.62, *P*=1.73×10^-10^), and *xerS* (β=-0.29, *P*=2.33×10^-09^), among others. The *atpD* gene is essential in Streptococcus and plays a role in energy metabolism, specifically in producing adenosine triphosphate (ATP) from adenosine diphosphate (ADP) (Song et al. 2005). Likewise, *xerS* encodes a protein that catalyzes DNA breakage and joining and may therefore play a role in recombination and, in turn, capsule switching. The RNA helicase unwinds RNA and modulates gene expression, similar to the ribosome-associated GTPase, which may influence the fitness of pneumococcal capsule switch mutants. Taken together, these findings suggest that although genomic variation in different pneumococcal backgrounds may affect the propensity to acquire novel capsule types into new genetic backgrounds across the phylogeny, the collective effect of the genetic background may influence capsule-switching propensity more than individual loci.

## Discussion

Despite the critical role of capsules in bacterial survival (3, 4), evasion of host immunity (6), and virulence (5), capsule-switching patterns remain poorly understood, even for clinically significant and well-studied bacterial pathogens. Here, we performed a comprehensive, large-scale phylogenomic analysis to quantify the frequency of capsule switching across capsule types and genetic backgrounds and to assess how genetic and ecological factors influence capsule-switching propensity in bacteria, as exemplified by the human-restricted respiratory pathogen pneumococcus. Remarkably, we show that pneumococcal strains expressing different capsule types acquire and lose capsules at unequal frequencies, depending on capsule type and genetic background. Interestingly, these capsule-switching patterns correlate with genetic and carriage-related capsule characteristics, including genetic similarity between switched cps loci, nasopharyngeal carriage duration, and capsule thickness, suggesting that genetic and ecological factors collectively influence capsule-switching dynamics in the pneumococcal population. These insights will enhance our understanding of the drivers of capsule switching, particularly the emergence and clonal expansion of novel vaccine-escape capsule types globally (Brueggemann et al. 2007; Moore et al. 2008; Croucher et al. 2011; Golubchik et al. 2012), especially in genetic backgrounds highly permissive to capsule switching.

The finding that some pneumococcal lineages and capsule types may be more permissive to capsule switching in the nasopharyngeal carriage population suggests the observed capsule-switching patterns are not entirely random (Joshi et al. 2020). Indeed, after normalizing the number of genomes included in the study for each capsule type, we identified several capsule types with a high propensity for capsule switching, including 18F, 18B, 35C, 29, and 33B. These capsule types did not include well-known, highly prevalent pediatric carriage and invasive disease capsule types (Johnson et al. 2010). In contrast, some pneumococcal genetic backgrounds express diverse capsule types, including those associated with serotype replacement in invasive disease after PCV introduction (Lo et al. 2022). The high variability in the number of capsule types expressed across lineages, as well as the number of lineages expressing individual capsule types, suggests that the propensity for capsule switching depends on both genetic background and capsule type. With the widespread introduction of PCVs, these capsule-switch-prone lineages and capsule types may pose major challenges for eradicating globally successful lineages (e.g., GPSC10) with capsule-specific vaccines, as these strains may readily switch to non-vaccine capsule types (Lo et al. 2022). These lineages may facilitate the emergence of novel vaccine-escape capsule types, thereby reducing the long-term population-level effectiveness of PCVs (Brueggemann et al. 2007; Wyres et al. 2013). While PCVs target the most highly switched capsule types, several non-vaccine capsule types also switch frequently. Interestingly, we have seen that prolonged use of effective PCVs against infrequently switched capsule types, such as 1 and 5 (Brueggemann et al. 2003; Løchen et al. 2022), significantly reduced the prevalence of the lineages harboring them, as these lineages are less likely to switch to other capsule types. Because novel pneumococcal capsule types continue to emerge and expand clonally, careful monitoring and investigation remain critical to predict and control capsule types likely to emerge, as well as their genetic backgrounds, to target them for elimination.

Several barriers are thought to influence homologous recombination and HGT in bacteria, including donor-recipient similarity (genetic), donor-recipient physical distance (ecological), and genomic adaptation of imported DNA (functional) barriers (Majewski and Cohan 1999; Popa et al. 2011; Popa and Dagan 2011). Our observation that only ∼4% of all possible capsule-type pairs directly switched exemplifies these barriers in pneumococcus. First, the pneumococcal capsule has also been hypothesized to act as a physical barrier to homologous recombination and HGT (76), suggesting that reduced encapsulation may promote pneumococcal diversification (Yother et al. 1986; Weiser and Kapoor 1999; Pearce et al. 2002; Chewapreecha et al. 2014). Ultimately, this hypothesis suggests that HGT of genomic loci, such as the *cps* cassette, leading to capsule switching, may be less common in highly encapsulated strains than in less encapsulated or unencapsulated strains(Ma et al. 2025). Asymmetry in pneumococcal capsule-switching frequency was evident across the phylogeny, possibly reflecting that transformation is not always bidirectional between the donor and recipient strains (Lattar et al. 2018). Our observation that pneumococcal strains expressing type 3 capsules readily replace other non-type 3 capsules in other strains but are rarely replaced by other capsule types supports the notion of asymmetry in capsule-switching frequency to and from specific capsule types. This directionality is plausible because thicker capsules typically persist longer during nasopharyngeal carriage (65), providing more opportunities for genetic exchange (Marks et al. 2012; Chaguza et al. 2016). However, the thick capsule itself might hinder the uptake of DNA (Ma et al. 2025), therefore, the high prevalence and persistence of type 3 strains during carriage isn’t necessarily driving those existing lineages to take on new non-type 3 *cps* loci, i.e., they are not behaving as recipients during colonization, but instead act as donors, spreading the type 3 capsule themselves far and wide onto many different genetic backgrounds.

In contrast, strains expressing other thin capsules may readily acquire naked exogenous type 3 *cps* loci DNA, leading to a higher switch from other capsule types to type 3 than vice versa. We also noted a few other capsule types, including 6D, 35C, 17F, 18A, 24F, 19B, 22F, and 10A, which were acquired substantially more often than they were replaced in the global pneumococcal phylogeny, suggesting that such potential hindering effects of the capsule may also be noticeable in other capsule types. However, we could not statistically assess the association between capsule acquisition and loss and capsule thickness because reliable capsule thickness data were unavailable for ∼90% of the capsule types described in this study. While previous studies have suggested that the capsule influences recombination and HGT (Chewapreecha et al. 2014; Haudiquet et al. 2024) and that unencapsulated strains may act as highways for these evolutionary processes leading to capsule switching (Chewapreecha et al. 2014; Andam and Hanage 2015; Bradshaw et al. 2020), our results did not reveal unexpectedly high capsule switching involving non-typeable (NT) and untypable pneumococcal strains compared to other capsule types. Because these unencapsulated strains rarely form clades, their unencapsulated states may be transient, and they may quickly revert to encapsulated versions, masking the effect of encapsulation on capsule acquisition in the present study. Taken together, our findings suggest that high bacterial encapsulation may impede exogenous DNA uptake and recombination, thereby decreasing the frequency of capsule switching to other capsule types and vice versa.

Other ecological and genetic factors, in addition to capsule thickness, also appear to influence capsule switching. Our regression analysis suggests that assayed and unassayed factors correlated with the duration of nasopharyngeal carriage strongly influence capsule switching. Because genetic exchange occurs exclusively during nasopharyngeal carriage (48), these results show the importance of prolonged carriage in increasing opportunities for capsule switching (Chaguza et al. 2016). Because strains expressing different capsule types colonize the nasopharynx for variable durations (58, 80), prolonged carriage may increase exposure and interactions between co-carried strains, thereby increasing opportunities for capsule switching. However, shorter carriage duration of certain capsule types may also imply faster transmission (to sustain them in the population), suggesting co-colonization in varied individuals, which itself may increase opportunities for capsule switching. However, the limited switching observed in the rarely carried capsule types, including 1 and 5, suggests that while other factors may also be at play, prolonged carriage duration may be more critical in influencing switching than increased transmissibility of rarely carried strains. In contrast to the observation that the capsule generally inhibits DNA uptake and transformation, we found that capsule switching for individual capsule types (such as type 3) occurs readily at the population level in strains expressing thicker capsules, unlike those with thinner capsules. Although this may seem counterintuitive, pneumococcal strains expressing thicker capsules are less susceptible to phagocytic killing (Weinberger et al. 2009), antibody-mediated killing (Abeyta et al. 2003), and mucus clearance (Nelson et al. 2007), which promotes carriage and increases the opportunities for capsule switching, albeit at a lower rate than for strains expressing thinner capsule types. Beyond these ecological factors, genetic factors may also influence capsule-switching frequency in pneumococci. Our findings show that the length of the *cps* locus cassette influences capsule switching, whereby smaller *cps* cassettes are more readily acquired than longer ones. These observations are consistent with the notion that the recombination of shorter DNA fragments occurs more frequently than that of longer fragments (Croucher et al. 2011; Apagyi et al. 2018). In addition, genetic similarity increases the frequency of capsule switching between capsule-type pairs. These findings align with previous reports that capsule switching occurs more frequently between similar cps cassettes, i.e., within a serogroup (30), again consistent with recombination rates increasing as sequence divergence decreases (Majewski et al. 2000).

Besides these capsule-specific genetic factors, inter-strain genomic variation in the rest of the pneumococcal background may also influence interactions between pneumococcal strains, potentially due to epistatic interactions between the *cps* and backbone (Croucher, Kagedan, et al. 2015; Joshi et al. 2020). Specifically, strongly outcompeting strains, possibly mediated by bacteriocins (Riley and Gordon 1999), may be co-carried less frequently, limiting DNA exchange between them. Others have also suggested that the inactivation of competence machinery genes may inhibit transformation and, consequently, bacterial diversification (Croucher et al. 2014). Our GWAS approach revealed several pneumococcal genomic variants statistically associated with capsule-switching propensity across lineages. Identifying only a few loci associated with capsule-switching propensity across genes with diverse functions suggests that the genetic background may collectively influence capsule switching rather than specific individual loci. While previous studies have suggested that some loci, including prophage insertions (Kwun et al. 2022) and other genomic changes (Durieux et al. 2019; Mazzamurro et al. 2024), influence bacterial transformation, our GWAS did not implicate these known loci. One possible explanation is that these previously reported loci may affect only a select few lineages, or that additional loci may also influence capsule switching. Nevertheless, our findings highlight the influence of the capsule itself, as well as the duration and prevalence of nasopharyngeal carriage, sequence similarity between *cps* loci, and the genetic background on the dynamics of pneumococcal capsule switching. However, follow-up GWAS and experimental studies are needed to identify and confirm additional loci influencing variability in transformation rates and the propensity for capsule switching in pneumococcal strains (Evans and Rozen 2013; Mazzamurro et al. 2024) beyond our preliminary proof-of-concept GWAS analysis.

In conclusion, our large-scale phylogenomics study provides comprehensive insights into the complex evolutionary dynamics of capsule switching in bacterial populations, as exemplified by the clinically significant human-restricted commensal and pathogen *S. pneumoniae.* We show how the propensity for capsule switching varies by capsule type, genetic background, and specific capsule-type pairs, and how factors associated with frequent or prolonged nasopharyngeal carriage influence it. Besides the carriage-related factors, other factors, including similarity and length of the switched *cps* loci, the capsule type itself and its properties e.g., thickness, the genetic background, and specific genomic loci, may influence the propensity for capsule switching, highlighting the role of genetic and ecological barriers to capsule switching, which require further investigation. Although computationally intensive, our analytical framework can be readily extended to investigate capsule switching in other encapsulated bacteria (Bellais et al. 2012; Wyres et al. 2015; Murray et al. 2023) and to study the evolutionary dynamics of other biological traits, such as antimicrobial resistance and virulence, beyond capsule switching. As shown in this study, collating large-scale datasets and applying our phylogenomics framework will provide novel insights into the drivers of phenotypic evolution in bacteria, facilitating prediction of the emergence of molecular traits, such as novel capsule types, across distinct bacterial genetic backgrounds. Such knowledge will ultimately inform disease prevention and control strategies, significantly tilting the scales in the human-bacteria arms race in favor of humans.

## Methods

### Global pneumococcal whole-genome sequencing dataset

We leveraged an extensive collection of pneumococcal genomes generated by the Global Pneumococcal Sequencing (GPS) project (https://www.pneumogen.net/gps/). We leveraged a whole-genome sequencing dataset comprising deposited sequence reads, draft genome assemblies, and the associated isolate metadata via the GPS Monocle database (https://data.monocle.sanger.ac.uk) [Accessed in November 2023]. The dataset used in the present analysis comprises sequences of carriage and invasive disease pneumococcal isolates from individuals of all ages across 56 countries and six continents **(Supplementary Data 1)**. We inferred capsule types from the isolates’ sequence reads using *in silico* serotyping methods implemented in SeroBA (version 1.0.0) (Epping et al. 2018). The sequencing data underwent quality control to assess for contamination as follows: 1) overall sequencing depth >20✕, 2) >60% reads mapping to *S. pneumoniae* using Kraken (version 2.1.2) (Wood and Salzberg 2014), 3) >60% mapping coverage of reference genome of the Pneumococcal Molecular Epidemiology Network (PMEN) global clone Spain23F-1 (GenBank accession: FM211187) (Croucher et al. 2009), 4) the total number of heterozygous single nucleotide polymorphisms (SNPs) <220 to exclude potential mixed samples, 5) <500 total number of contigs, and 6) total length of the assembled genome ranging between 1.9–2.3 Mb (Tettelin et al. 2001).

We identified pneumococcal sequence types (STs), or clones, using the multilocus sequence typing (MLST) approach (90) with the pneumococcal MLST scheme (91) and the “mlst” script (version 2.22.0) (https://github.com/tseemann/mlst). Because the MLST scheme assigns clones based on only seven housekeeping genes out of several thousand genes in the pneumococcal genome, sometimes it may have low discriminatory power, as highly related strains may be assigned different clone names. Furthermore, some analyses require broader groupings of pneumococcal isolates into clusters, lineages, or genetic backgrounds with consistent nomenclature, such as clonal complexes that group closely related STs (Enright and Spratt 1998). Since the clonal complexes lack resolution, as they are defined based on a few genomic loci in the pneumococcal genome, we therefore assigned consistent pneumococcal lineages or genetic backgrounds based on the global pneumococcal sequence cluster (GPSC) (Gladstone et al. 2019) implemented in the POPulation Partitioning Using Nucleotide K-mers (PopPUNK) framework (Lees et al. 2019).

We downsampled the 21,155 draft-assembled genomes to 10,558 to make the computationally intensive downstream genomic analyses feasible. Based on our previous analysis of the pneumococcal genomes from the GPS project (Gladstone et al. 2019), we postulated that the dataset of ∼10,000 genomes was feasible for this study. When sampling the genomes, we randomly selected up to five genomes for each combination of capsule type, genetic background or lineage (STs and GPSCs), and country. This approach enabled the inclusion of rare capsule types and genetic backgrounds while simultaneously capturing the geographical diversity of the genomes in the final dataset and analysis. Compared with random selection, our sampling approach included rare capsule types and genetic backgrounds and avoided overrepresenting highly prevalent capsule types that are oversampled in certain countries, which contributed more samples for whole-genome sequencing.

### Comparative genomic analysis

Before downstream phylogenetic and pan-genome analysis, we assembled the pneumococcal genomes using SPAdes (version 3.16.4) (Bankevich et al. 2012). All selected genomes passed quality checks based on a previously described approach (Gladstone et al. 2019) that involved genomic DNA extracted from fresh, pure overnight cultures of single colonies using the previously described methods. To generate a reference-based whole-genome alignment of the 10,558 selected genomes, we mapped sequence reads of each isolate against the ATCC 700669 complete pneumococcal reference genome (GenBank accession: NC_011900.1) using Snippy (version 4.6.0) (https://github.com/tseemann/snippy). We also included a *Streptococcus mitis* genome (GenBank accession: CP067992.1) when generating the whole-genome alignment as an outgroup species. We ran Snippy with default values (minimum mapping quality of 60, minimum base quality of 13, and minimum site depth of 13 for variant calling; maximum allowed value of 10 for soft clipping to generate and align consensus nucleotide sequences). We used a reference-based approach to align the genomes because of the large dataset size and our interest in core-genome changes for downstream analysis, including phylogenetic reconstruction. We inferred these core-genome nucleotide changes from positions in the multiple sequence alignment that contained unambiguous nucleotide data for all isolates.

### Phylogenetic reconstruction

We generated an approximate maximum-likelihood phylogenetic tree using IQ-TREE (version 2.0.3) (Nguyen et al. 2015). We ran IQ-TREE on a shorter alignment of polymorphic sites generated from the initial alignment using SNP-sites (version 2.5.1) (95) with the “-c” option to include only variable positions with unambiguous nucleotides, which capture variation in the pneumococcal core genome. We rooted the phylogenetic tree using the *Streptococcus mitis* genome included in the generated whole-genome alignment. We processed the phylogenetic tree, including excluding the reference genome, outgroup-based rooting, and visualization, using ape (version 5.7.1) (Paradis et al. 2004) and phytools (version 2.1.1) (Revell 2012). Compared to the pan-genome alignment approach, the reference-based alignment method includes variable sites in aligned non-coding regions.

Because of the large dataset size, we did not perform a recombination analysis because it is computationally intensive and no robust tools exist to accurately detect and remove recombination events in species-wide, multi-clonal bacterial phylogenies, where divergence signals may reflect processes beyond recombination, such as geographical separation. Most widely used tools for detecting and removing recombination events in bacterial genomes, such as Gubbins (Croucher, Page, et al. 2015), are suitable for clonal populations; therefore, we did not use them in the present study. However, previous studies suggest that the topology of phylogenetic trees inferred from whole-genome alignments is usually robust to recombination (Hedge and Wilson 2014).

### Quantifying capsule switching patterns of specific capsule types and genetic backgrounds

We quantified the number of unique genetic backgrounds or lineages expressing a particular capsule type to assess the propensity for capsule switching across capsule types, expecting that frequently switched capsule types would be found in more lineages than less frequently switched ones. On the other hand, we quantified the propensity, or permissiveness, of a particular genetic background to capsule switching by the number of capsule types expressed in the lineage. Consequently, we considered genetic backgrounds with more capsule types more permissive to switching than those with fewer. We adjusted estimates of the propensity of specific capsule types and genetic backgrounds to capsule switching by dividing each estimate by the number of genomes included in the analysis and multiplying by 1,000 to obtain estimates per 1,000 sampled genomes.

Using the number of unique backgrounds expressing each capsule type to quantify the propensity of capsule types to undergo switching may underestimate this propensity, as some capsule types are acquired multiple times on different phylogenetic branches within the same genetic background. To address this, we employed discrete stochastic ancestral state reconstruction to estimate the acquisition and loss frequencies of each capsule type across the global pneumococcal phylogeny. By accounting for the phylogeny, this approach provides more accurate estimates of capsule acquisition and loss frequencies for each capsule type, while accounting for multiple independent acquisitions and losses. We defined two discrete states, i.e., “0” and “1”, for each capsule type, corresponding to its presence or absence in all genomes in the phylogeny. We defined the loss of a capsule type as any switch from one capsule type to another, including potential deletion of the cps locus, which renders a subset of strains untypable. We fitted a continuous-time reversible Markov model for capsule-type evolution, then simulated 100 stochastic character histories using the “make.simmap” function in phytools (version 2.1.1) (Huelsenbeck et al. 2003; Bollback 2006; Revell 2012). We plotted the resulting posterior density of stochastic mapping on the global pneumococcal phylogeny using the “densityMap” function in phytools (version 2.1.1) (Huelsenbeck et al. 2003; Bollback 2006; Revell 2012). We summarized the results of the stochastic character mapping to estimate the number of changes between states in the state transition matrix, i.e., “0→1” reflecting capsule acquisitions and “1→0” reflecting capsule replacement or switch to another capsule type, using the “describe.simmap” function in phytools (version 2.1.1) (Huelsenbeck et al. 2003; Bollback 2006; Revell 2012). We adjusted capsule acquisition and loss frequencies by dividing each estimate by the number of genomes included in the analysis and multiplying by 1,000 to obtain acquisitions or losses per 1,000 sampled genomes.

To quantify the frequency of capsule switching between specific pairs of capsule types, we repeated the ancestral state reconstruction, this time defining three discrete states corresponding to the first serotype in the pair (state 1), the second capsule type in the pair (state 2), and all other capsules together (state 3). In this case, inferred direct capsule switches between specific capsule types corresponded to the “1→2” and “2→1” changes in the inferred state transition matrix. The “3→1” and “3→2”, “1→3”, and “2→3” transitions correspond to capsule switches to and from the specific pair of capsule types. We summarized the results similarly, as described in the previous paragraph. We adjusted capsule-switch frequencies by dividing each estimate by the number of genomes included in the analysis and multiplying by 1,000 to obtain acquisitions or losses per 1,000 sampled genomes.

### Association between genetic, ecological, and epidemiological covariates correlates with capsule switching

We used Poisson generalized linear regression models to assess the association between the propensity of capsule types to undergo capsule switching and several genetic and ecological factors, such as nasopharyngeal carriage duration, as independent covariates. First, we evaluated the association between capsule acquisition frequency and nasopharyngeal carriage duration, *cps* locus size, and median capsule thickness. Using the ancestral state reconstruction approach described in the previous section, we inferred capsule acquisition frequency as the number of transitions (switches) from other capsule types to the capsule type of interest. The carriage duration and capsule thickness data were obtained from previous studies (Weinberger et al. 2009; Chaguza, Senghore, et al. 2020), while the *cps* locus size was based on the data published by Bentley et al (Bentley et al. 2006) and is available as part of the SeroBA (Epping et al. 2018) database for *in silico* pneumococcal serotyping (https://github.com/sanger-pathogens/seroba/blob/master/database/). Pneumococcal epidemiological studies primarily focus on invasive disease rather than asymptomatic nasopharyngeal carriage. In addition, serotypes are typically determined using laborious traditional serotyping methods, such as the Quellung reaction, latex agglutination, and multiplex PCR (Pai et al. 2006), and reliable data are limited or nonexistent for most of the capsule types analyzed in this study. The lack of extensive carriage data largely reflects the primary focus of traditional methods on a subset of serotypes included in pneumococcal conjugate vaccines (PCV). In contrast, less laborious, genome-based serotyping approaches can reliably infer nearly all serotypes with high sensitivity and specificity (Epping et al. 2018).

We formulated three regression models as follows. Model #1: □□□(□_□_) = *□_1_□ _1□_ + □□□□□□*(□_□_), Model #2: □□□(□_□_) = *□_1_□_1□_ + □_2_□_2_*_□_ _ □□□□□□(□_□_), and Model #3: □□□(□_□_) = □*_1_□_1□_ + □_2_□_2□_ + □_3_□_3□_* + □□□□□□(□_□_), where □_□_ represents the capsule acquisition frequency (count) for the ith strain or genome, represents the number of genomes included in the analysis per capsule type,□*_1□_* represents the carriage duration, □*_2□_*the *cps* locus size, and □*_3□_* the median capsule thickness, while *□_1_, □_2_*, and □_3_ represent the coefficients for the carriage duration, *cps* locus size, and capsule thickness. The term□_□_ represents the Poisson parameter, which represents the number of occurrences (e.g., capsule acquisitions or losses) per unit time. To compare the regression coefficients, we adjusted for each covariate included in the models. We included an offset term in each model to adjust for the number of genomes included in the analysis for each capsule type, thereby minimizing sampling bias by capsule type. We repeated the regression with similarly formulated models using a different outcome: capsule replacement frequency. We defined capsule replacement frequency as the number of capsule switches from a specific capsule type to any other capsule type using the ancestral state reconstruction approach described in the previous section. Next, we fitted these similarly structured models with the capsule acquisition-to-loss ratio (A/L ratio) as the outcome. We fitted a Gaussian generalized regression model. Model #1:□□ _ □_1_□_1□_ + □□□□□□(□_□_), Model #2: □□ _ □_1_□_1_□ _ □_2_□_2□_ + □□□□□□(□_□_) + □_□_, and Model #3: □_□_ = *□_1_□ _1_□ + □_2_□_2_□ + □_3_□_3_□ _ □□□□□*□(□_□_) + □_□_. Here □_□_ is the A/L ratio, and □_□_ represents the residual standard error.

To assess the effect of the genetic and ecological factors on the frequency of capsule switching between specific pairs of capsule types, we also performed a Poisson regression analysis with two models. Model #1:□□□(□_□_)_ = *□_1_□_1□_ + □_2_□_2□_ + □□□□□□*(□_□_)and model #2:□□□(□_□_) = *□_1_□_1□_ + □_2_□_2□_ + □_3_□_3□_ + □_4_□_4□_* + □□□□□□(□_□_) In each model □_□_represents the frequency (count) of capsule switching between ith capsule type pair,□_□_ represents the number of genomes included in the analysis per capsule type,□_1□_ represents the pairwise *cps* loci genetic similarity,□_2□_ represents the difference in *cps* locus sizes, □_3□_ represents the carriage duration of the replaced capsule type, and□_3□_ represents the carriage duration of the replacing capsule type, while *□_1_, □_2_, □_3_*, and □*_4_* represent the coefficients for the four covariates. Similarly, represents the Poisson parameter, which is the number of occurrences (e.g., capsule acquisitions or losses) per unit time. We also adjusted each covariate for easier comparison and included an offset term in each model to adjust for the number of genomes. Here, we specified the offset term as the log-transformed product of the number of genomes included in this study for the replaced and replacing capsule type for each pair of capsule types. We generated the covariate representing pairwise genetic similarity between cps locus sequences using the “dist” command in Mash (version 2.3) (Ondov et al. 2016). We used reference *cps* sequences from a database used by SeroBA (86) for in silico pneumococcal serotyping (https://github.com/sanger-pathogens/seroba/blob/master/database/), including the initial set of *cps* sequences reported by Bentley et al. (Bentley et al. 2006) and those for newly discovered capsule types.

### Assessing genetic factors associated with the frequency of capsule switching across different genetic backgrounds

We conducted a bacterial genome-wide association study (GWAS) to investigate the relationship between pneumococcal genomic variation and the propensity of pneumococcal genetic backgrounds or lineages to undergo switching. We defined each lineage’s propensity for capsule switching as the adjusted number of unique capsule types expressed per 1,000 sampled genomes, controlling for sampling bias, since oversampled lineages may harbor many capsule types. We based genomic variation in randomly selected genomes from each lineage on unitig sequences or variable-length k-mer sequences generated from each isolate’s draft assembly. We selected up to five genomes per lineage from lineages with at least ten sequenced genomes. We used these unitigs to effectively capture genomic variation at nucleotide substitution, insertion, and deletion levels within the conserved (core) and non-conserved (accessory) genomes. We chose this approach because it is particularly suitable for pneumococcus, given its highly recombinogenic nature (Croucher et al. 2011; Croucher et al. 2013; Chewapreecha et al. 2014; Chaguza et al. 2016). We performed the GWAS analysis using linear mixed-effects models implemented in Pyseer (version 1.3.7-dev) (Lees et al. 2018). These linear mixed models robustly control for bacterial population structure, which can confound GWAS analyses (Power et al. 2017). Specifically, we generated and specified a kinship matrix, which defined the variances and covariances of the random effects for each genome using the “similarity_pyseer” script in Pyseer to account for population structure in the GWAS. We excluded essentially rare and universally present unitigs with allele frequencies <5% and >95%, respectively. We inferred unitigs statistically associated with capsule-switching propensity when the adjusted statistical significance for each variant was <0.05, correcting for multiple testing using the Bonferroni method. The number of unitigs used for the Bonferroni adjustment equaled the total number of unique presence and absence patterns in the dataset, inferred using a Pyseer script (https://github.com/mgalardini/pyseer/blob/master/scripts/count_patterns.py). Based on this, we defined the genome-wide statistical significance threshold as *P* <1.20×10^-07^, i.e., /G, where =0.05 and G=415,849 is the number of unique presence and absence patterns of the unitigs. We assessed potential issues with the GWAS analysis, possibly arising from problems with correcting for population structure, by examining the GWAS output with quantile-quantile (QQ) plots using qqman (version 0.1.7) (Turner). We annotated unitig sequences statistically associated with capsule-switching propensity in the genetic backgrounds using BLASTN (version 2.5.0+) (Altschul et al. 1990) and BioPython (Cock et al. 2009). We compared each unitig sequence against the ATCC 700669 pneumococcal reference genome (GenBank accession: NC_011900) to identify genomic features found in the genomic regions where the unitigs mapped. We summarized and visualized the GWAS results using Manhattan plots in R (version 4.0.3) [https://www.R-project.org/].

## Data availability

We have deposited the whole-genome sequencing data for the study isolates in the European Nucleotide Archive (ENA) and provided the accession numbers and other isolate metadata in **Supplementary Data 1**. We describe all other data supporting the findings of this study in this paper or in the supplementary material. We have also provided additional raw data on GitHub: https://github.com/ChrispinChaguza/PneumococcalCapsuleSwitchGlobal.

## Code availability

We describe all tools and methods used for the analysis in the Materials and Methods sections. Other scripts used in this analysis are available publicly on GitHub: https://github.com/ChrispinChaguza/PneumococcalCapsuleSwitchGlobal.

## Acknowledgments

We thank the Global Pneumococcal Sequencing (GPS) project partners for collecting, sequencing, and publicly sharing the data for the pneumococcal samples. D.M.W. is supported by R01AI195985 from the NIH/NIAID. C.P.A. is supported by a grant from the National Institutes of Health (Award no. R35GM142924). The Wellcome Sanger Institute activities were broadly supported by the Gates Foundation (grant code OPP1034556) and Wellcome Trust (Award numbers: 098051 and 206194). R.S.H. acknowledges funding from the Department of Health’s National Institute for Health and Care Research (NIHR) Biomedical Research Centre at University College London and University College London Hospitals NHS Foundation Trust (UCLH). R.S.H. is a NIHR Senior Investigator. We acknowledge the Pathogen Genomics team at the Wellcome Sanger Institute and the Yale Center for Research Computing (YCRC) for providing high-performance computing infrastructure. We also thank members of the Weinberger and Pitzer research labs for feedback on this work.

## Author contributions

C.C. and D.M.W. conceived the study; members of the GPS consortium (https://www.pneumogen.net/gps/) collected the original samples and clinical data and performed microbiological and molecular sample processing; S.W.L., R.A.G., and S.D.B. oversaw whole genome sequencing and data processing; C.C. performed the bioinformatics and statistical analyses with support from D.M.W.; D.M.W. supervised the study; A.Y., C.P.A., and D.M.W. contributed to the discussions of the data analysis and interpretation. C.C. and D.M.W. drafted the manuscript; D.M.W., L.M-G, R.F.B., and S.D.B. secured funds and supervised the project; All authors reviewed and approved the manuscript.

## ^†^Global Pneumococcal Sequencing (GPS) consortium

Alejandra Corso, Paula Gagetti, Abdullah W. Brooks, Md Hasanuzzaman, Samir K. Saha, Senjuti Saha, Alexander Davydov, Leonid Titov, Samanta Cristine Grassi Almeida, Paul Turner, Chunjiang Zhao, Hui Wang, Margaret Ip, Pak Leung Ho, Pierra Law, Jeremy D. Keenan, Robert Cohen, Emmanuelle Varon, Eric Sampane-Donkor, Balaji Veeraraghavan, Geetha Nagaraj, K. L. Ravikumar, J. Yuvaraj, Varun Shamanna Noga, Rachel Benisty, Ron Dagan, Godfrey Bigogo, Jennifer Verani, Anmol Kiran, Dean B. Everett, Jennifer Cornick, Maaike Alaerts, Shamala Devi Sekaran, Stuart C. Clarke, Benild Moiane, Betuel Sigauque, Helio Mucavele, Andrew J. Pollard, Rama Kandasamy, Philip E. Carter, Stephen K. Obaro, Deborah Lehmann, Rebecca Ford, Theresa J. Ochoa, Anna Skoczynska, Ewa Sadowy, Waleria Hryniewicz, Weronika Puzia, Sanjay Doiphode, Ekaterina Egorova, Elena Voropaeva, Yulia Urban, Tamara Kastrin, Kedibone Ndlangisa, Linda De Gouveia, Mushal Ali, Nicole Wolter, Cebile Lekhuleni, Carmen Muñoz Almagro, Alba Redin Alonso, Desiree Henares, Somporn Srifuengfung, Martin Antonio, Brenda Kwambana-Adams, Ebenezer Foster-Nyarko, Ebrima Bojang, Peggy-Estelle Tientcheu, Jennifer Moïsi, Michele Nurse-Lucas, Patrick E. Akpaka, Özgen Köseoglu Eser, Anne von Gottberg, Mignon du Plessis, Shabir A. Madhi, Anthony Scott, David Aanensen, John A. Lees, Rebecca A. Gladstone, Gerry Tonkin-Hill, Chrispin Chaguza, David Cleary, Kate Mellor, Sophie Belman, Bernard Beall, Gail Rodgers, Paulina A. Hawkins, Anne J. Blaschke, Robert F. Breiman, and Nicole L. Pershing.

## References

Abeyta M, Hardy GG, Yother J. 2003. Genetic alteration of capsule type but not PspA type affects accessibility of surface-bound complement and surface antigens of Streptococcus pneumoniae. Infect Immun [Internet] 71:218–225. Available from: 10.1128/IAI.71.1.218-225.2003

Alcalá B, Arreaza L, Salcedo C, Uría MJ, De La Fuente L, Vázquez JA. 2002. Capsule switching among C:2b:P1.2,5 meningococcal epidemic strains after mass immunization campaign, Spain. Emerg. Infect. Dis. [Internet] 8:1512–1514. Available from: 10.3201/eid0812.020081

Altschul SF, Gish W, Miller W, Myers EW, Lipman DJ. 1990. Basic local alignment search tool. J. Mol. Biol. [Internet] 215:403–410. Available from: 10.1016/S0022-2836(05)80360-2

Andam CP, Hanage WP. 2015. Mechanisms of genome evolution of Streptococcus. Infect. Genet. Evol. [Internet] 33:334–342. Available from: 10.1016/j.meegid.2014.11.007

Apagyi KJ, Fraser C, Croucher NJ. 2018. Transformation Asymmetry and the Evolution of the Bacterial Accessory Genome. Mol Biol Evol [Internet] 35:575–581. Available from: 10.1093/molbev/msx309

Arnold BJ, Huang I-T, Hanage WP. 2022. Horizontal gene transfer and adaptive evolution in bacteria. Nat. Rev. Microbiol. [Internet] 20:206–218. Available from: 10.1038/s41579-021-00650-4

Bankevich A, Nurk S, Antipov D, Gurevich AA, Dvorkin M, Kulikov AS, Lesin VM, Nikolenko SI, Pham S, Prjibelski AD, et al. 2012. SPAdes: a new genome assembly algorithm and its applications to single-cell sequencing. J. Comput. Biol. [Internet] 19:455– 477. Available from: 10.1089/cmb.2012.0021

Bellais S, Six A, Fouet A, Longo M, Dmytruk N, Glaser P, Trieu-Cuot P, Poyart C. 2012. Capsular switching in group B Streptococcus CC17 hypervirulent clone: a future challenge for polysaccharide vaccine development. J. Infect. Dis. [Internet] 206:1745–1752. Available from: 10.1093/infdis/jis605

Bentley SD, Aanensen DM, Mavroidi A, Saunders D, Rabbinowitsch E, Collins M, Donohoe K, Harris D, Murphy L, Quail MA, et al. 2006. Genetic analysis of the capsular biosynthetic locus from all 90 pneumococcal serotypes. PLoS Genet. [Internet] 2:e31. Available from: 10.1371/journal.pgen.0020031

Blackwell GA, Hunt M, Malone KM, Lima L, Horesh G, Alako BTF, Thomson NR, Iqbal Z. 2021. Exploring bacterial diversity via a curated and searchable snapshot of archived DNA sequences. PLoS Biol. [Internet] 19:e3001421. Available from: 10.1371/journal.pbio.3001421

Bollback JP. 2006. SIMMAP: stochastic character mapping of discrete traits on phylogenies. BMC Bioinformatics [Internet] 7:88. Available from: 10.1186/1471-2105-7-88

Bradshaw JL, Rafiqullah IM, Robinson DA, McDaniel LS. 2020. Transformation of nonencapsulated Streptococcus pneumoniae during systemic infection. Sci Rep [Internet] 10:18932. Available from: 10.1038/s41598-020-75988-5

Brueggemann AB, Griffiths DT, Meats E, Peto T, Crook DW, Spratt BG. 2003. Clonal relationships between invasive and carriage Streptococcus pneumoniae and serotype-and clone-specific differences in invasive disease potential. J. Infect. Dis. [Internet] 187:1424– 1432. Available from: 10.1086/374624

Brueggemann AB, Pai R, Crook DW, Beall B. 2007. Vaccine escape recombinants emerge after pneumococcal vaccination in the United States. PLoS Pathog. [Internet] 3:e168. Available from: 10.1371/journal.ppat.0030168

Chaguza C, Andam CP, Harris SR, Cornick JE, Yang M, Bricio-Moreno L, Kamng’ona AW, Parkhill J, French N, Heyderman RS, et al. 2016. Recombination in Streptococcus pneumoniae Lineages Increase with Carriage Duration and Size of the Polysaccharide Capsule. MBio [Internet] 7. Available from: 10.1128/mBio.01053-16

Chaguza C, Cornick JE, Andam CP, Gladstone RA, Alaerts M, Musicha P, Peno C, Bar-Zeev N, Kamng’ona AW, Kiran AM, et al. 2017. Population genetic structure, antibiotic resistance, capsule switching and evolution of invasive pneumococci before conjugate vaccination in Malawi. Vaccine [Internet] 35:4594–4602. Available from: 10.1016/j.vaccine.2017.07.009

Chaguza C, Heinsbroek E, Gladstone RA, Tafatatha T, Alaerts M, Peno C, Cornick JE, Musicha P, Bar-Zeev N, Kamng’ona A, et al. 2020. Early Signals of Vaccine-driven Perturbation Seen in Pneumococcal Carriage Population Genomic Data. Clin. Infect. Dis. [Internet] 70:1294–1303. Available from: 10.1093/cid/ciz404

Chaguza C, Senghore M, Bojang E, Lo SW, Ebruke C, Gladstone RA, Tientcheu P-E, Bancroft RE, Worwui A, Foster-Nyarko E, et al. 2020. Carriage Dynamics of Pneumococcal Serotypes in Naturally Colonized Infants in a Rural African Setting During the First Year of Life. Front Pediatr [Internet] 8:587730. Available from: 10.3389/fped.2020.587730

Chewapreecha C, Harris SR, Croucher NJ, Turner C, Marttinen P, Cheng L, Pessia A, Aanensen DM, Mather AE, Page AJ, et al. 2014. Dense genomic sampling identifies highways of pneumococcal recombination. Nat. Genet. [Internet] 46:305–309. Available from: 10.1038/ng.2895

Cock PJA, Antao T, Chang JT, Chapman BA, Cox CJ, Dalke A, Friedberg I, Hamelryck T, Kauff F, Wilczynski B, et al. 2009. Biopython: freely available Python tools for computational molecular biology and bioinformatics. Bioinformatics [Internet] 25:1422– 1423. Available from: 10.1093/bioinformatics/btp163

Coffey TJ, Enright MC, Daniels M, Morona JK, Morona R, Hryniewicz W, Paton JC, Spratt BG. 1998. Recombinational exchanges at the capsular polysaccharide biosynthetic locus lead to frequent serotype changes among natural isolates of Streptococcus pneumoniae. Mol Microbiol [Internet] 27:73–83. Available from: 10.1046/j.1365-2958.1998.00658.x

Cowley LA, Petersen FC, Junges R, Jimson D Jimenez M, Morrison DA, Hanage WP. 2018. Evolution via recombination: Cell-to-cell contact facilitates larger recombination events in Streptococcus pneumoniae. PLoS Genet. [Internet] 14:e1007410. Available from: 10.1371/journal.pgen.1007410

Cress BF, Englaender JA, He W, Kasper D, Linhardt RJ, Koffas MAG. 2014. Masquerading microbial pathogens: capsular polysaccharides mimic host-tissue molecules. FEMS Microbiol. Rev. [Internet] 38:660–697. Available from: 10.1111/1574-6976.12056

Croucher NJ, Finkelstein JA, Pelton SI, Mitchell PK, Lee GM, Parkhill J, Bentley SD, Hanage WP, Lipsitch M. 2013. Population genomics of post-vaccine changes in pneumococcal epidemiology. Nat. Genet. [Internet] 45:656–663. Available from: 10.1038/ng.2625

Croucher NJ, Hanage WP, Harris SR, McGee L, van der Linden M, de Lencastre H, Sá-Leão R, Song J-H, Ko KS, Beall B, et al. 2014. Variable recombination dynamics during the emergence, transmission and “disarming” of a multidrug-resistant pneumococcal clone. BMC Biol [Internet] 12:49. Available from: 10.1186/1741-7007-12-49

Croucher NJ, Harris SR, Fraser C, Quail MA, Burton J, van der Linden M, McGee L, von Gottberg A, Song JH, Ko KS, et al. 2011. Rapid pneumococcal evolution in response to clinical interventions. Science [Internet] 331:430–434. Available from: 10.1126/science.1198545

Croucher NJ, Kagedan L, Thompson CM, Parkhill J, Bentley SD, Finkelstein JA, Lipsitch M, Hanage WP. 2015. Selective and genetic constraints on pneumococcal serotype switching. PLoS Genet. [Internet] 11:e1005095. Available from: 10.1371/journal.pgen.1005095

Croucher NJ, Page AJ, Connor TR, Delaney AJ, Keane JA, Bentley SD, Parkhill J, Harris SR. 2015. Rapid phylogenetic analysis of large samples of recombinant bacterial whole genome sequences using Gubbins. Nucleic Acids Res. [Internet] 43:e15. Available from: 10.1093/nar/gku1196

Croucher NJ, Walker D, Romero P, Lennard N, Paterson GK, Bason NC, Mitchell AM, Quail MA, Andrew PW, Parkhill J, et al. 2009. Role of conjugative elements in the evolution of the multidrug-resistant pandemic clone Streptococcus pneumoniaeSpain23F ST81. J. Bacteriol. [Internet] 191:1480–1489. Available from: 10.1128/JB.01343-08

Durieux I, Ginevra C, Attaiech L, Picq K, Juan P-A, Jarraud S, Charpentier X. 2019. Diverse conjugative elements silence natural transformation in species. Proc Natl Acad Sci U S A [Internet] 116:18613–18618. Available from: 10.1073/pnas.1909374116

Enright MC, Spratt BG. 1998. A multilocus sequence typing scheme for Streptococcus pneumoniae: identification of clones associated with serious invasive disease. Microbiology [Internet] 144 (Pt 11):3049–3060. Available from: 10.1099/00221287-144-11-3049

Epping L, van Tonder AJ, Gladstone RA, The Global Pneumococcal Sequencing Consortium, Bentley SD, Page AJ, Keane JA. 2018. SeroBA: rapid high-throughput serotyping of Streptococcus pneumoniae from whole genome sequence data. Microb Genom [Internet] 4. Available from: 10.1099/mgen.0.000186

Evans BA, Rozen DE. 2013. Significant variation in transformation frequency in Streptococcus pneumoniae. ISME J [Internet] 7:791–799. Available from: 10.1038/ismej.2012.170

Gladstone RA, Jefferies JM, Tocheva AS, Beard KR, Garley D, Chong WW, Bentley SD, Faust SN, Clarke SC. 2015. Five winters of pneumococcal serotype replacement in UK carriage following PCV introduction. Vaccine [Internet] 33:2015–2021. Available from: 10.1016/j.vaccine.2015.03.012

Gladstone RA, Lo SW, Lees JA, Croucher NJ, van Tonder AJ, Corander J, Page AJ, Marttinen P, Bentley LJ, Ochoa TJ, et al. 2019. International genomic definition of pneumococcal lineages, to contextualise disease, antibiotic resistance and vaccine impact. EBioMedicine [Internet] 43:338–346. Available from: 10.1016/j.ebiom.2019.04.021

Goldblatt D. 2000. Conjugate vaccines. Clin. Exp. Immunol. [Internet] 119:1–3. Available from: 10.1046/j.1365-2249.2000.01109.x

Golden AR, Adam HJ, Gilmour MW, Baxter MR, Martin I, Nichol KA, Demczuk WHB, Hoban DJ, Zhanel GG. 2015. Assessment of multidrug resistance, clonality and virulence in non-PCV-13 Streptococcus pneumoniae serotypes in Canada, 2011-13. J. Antimicrob. Chemother. [Internet] 70:1960–1964. Available from: 10.1093/jac/dkv061

Golubchik T, Brueggemann AB, Street T, Gertz RE Jr, Spencer CCA, Ho T, Giannoulatou E, Link-Gelles R, Harding RM, Beall B, et al. 2012. Pneumococcal genome sequencing tracks a vaccine escape variant formed through a multi-fragment recombination event. Nat Genet [Internet] 44:352–355. Available from: 10.1038/ng.1072

Hamaguchi S, Zafar MA, Cammer M, Weiser JN. 2018. Capsule Prolongs Survival of Streptococcus pneumoniae during Starvation. Infect. Immun. [Internet] 86. Available from: 10.1128/IAI.00802-17

Hathaway LJ, Grandgirard D, Valente LG, Täuber MG, Leib SL. 2016. Streptococcus pneumoniae capsule determines disease severity in experimental pneumococcal meningitis. Open Biol. [Internet] 6. Available from: 10.1098/rsob.150269

Haudiquet M, Le Bris J, Nucci A, Bonnin RA, Domingo-Calap P, Rocha EPC, Rendueles O. 2024. Capsules and their traits shape phage susceptibility and plasmid conjugation efficiency. Nat. Commun. [Internet] 15:2032. Available from: 10.1038/s41467-024-46147-5

Hedge J, Wilson DJ. 2014. Bacterial phylogenetic reconstruction from whole genomes is robust to recombination but demographic inference is not. MBio [Internet] 5:e02158. Available from: 10.1128/mBio.02158-14

Huelsenbeck JP, Nielsen R, Bollback JP. 2003. Stochastic mapping of morphological characters. Syst. Biol. [Internet] 52:131–158. Available from: 10.1080/10635150390192780

Hyams C, Camberlein E, Cohen JM, Bax K, Brown JS. 2010. The Streptococcus pneumoniae capsule inhibits complement activity and neutrophil phagocytosis by multiple mechanisms. Infect. Immun. [Internet] 78:704–715. Available from: 10.1128/IAI.00881-09

Jefferies JMC, Smith A, Clarke SC, Dowson C, Mitchell TJ. 2004. Genetic analysis of diverse disease-causing pneumococci indicates high levels of diversity within serotypes and capsule switching. J. Clin. Microbiol. [Internet] 42:5681–5688. Available from: 10.1128/JCM.42.12.5681-5688.2004

Johnson HL, Deloria-Knoll M, Levine OS, Stoszek SK, Freimanis Hance L, Reithinger R, Muenz LR, O’Brien KL. 2010. Systematic evaluation of serotypes causing invasive pneumococcal disease among children under five: the pneumococcal global serotype project. PLoS Med. [Internet] 7. Available from: 10.1371/journal.pmed.1000348

Joshi SS, Al-Mamun MA, Weinberger DM. 2020. Correlates of nonrandom patterns of serotype switching in pneumococcus. J. Infect. Dis. [Internet] 221:1669–1676. Available from: 10.1093/infdis/jiz687

Kadioglu A, Weiser JN, Paton JC, Andrew PW. 2008. The role of Streptococcus pneumoniae virulence factors in host respiratory colonization and disease. Nat Rev Microbiol [Internet] 6:288–301. Available from: 10.1038/nrmicro1871

Khan UB, Jauneikaite E, Andrews R, Chalker VJ, Spiller OB. 2022. Identifying large-scale recombination and capsular switching events in strains causing disease in adults in the UK between 2014 and 2015. Microb Genom [Internet] 8. Available from: 10.1099/mgen.0.000783

King SJ, Leigh JA, Heath PJ, Luque I, Tarradas C, Dowson CG, Whatmore AM. 2002. Development of a multilocus sequence typing scheme for the pig pathogen Streptococcus suis: identification of virulent clones and potential capsular serotype exchange. J. Clin. Microbiol. [Internet] 40:3671–3680. Available from: 10.1128/JCM.40.10.3671-3680.2002

Kwun MJ, Ion AV, Cheng H-C, D’Aeth JC, Dougan S, Oggioni MR, Goulding DA, Bentley SD, Croucher NJ. 2022. Post-vaccine epidemiology of serotype 3 pneumococci identifies transformation inhibition through prophage-driven alteration of a non-coding RNA. Genome Med [Internet] 14:144. Available from: 10.1186/s13073-022-01147-2

Lancellotti M, Guiyoule A, Ruckly C, Hong E, Alonso J-M, Taha M-K. 2006. Conserved virulence of C to B capsule switched Neisseria meningitidis clinical isolates belonging to ET-37/ST-11 clonal complex. Microbes Infect. [Internet] 8:191–196. Available from: 10.1016/j.micinf.2005.06.012

Lattar SM, Wu X, Brophy J, Sakai F, Klugman KP, Vidal JE. 2018. A Mechanism of Unidirectional Transformation, Leading to Antibiotic Resistance, Occurs within Nasopharyngeal Pneumococcal Biofilm Consortia. mBio [Internet] 9. Available from: 10.1128/mBio.00561-18

Lees JA, Galardini M, Bentley SD, Weiser JN, Corander J. 2018. pyseer: a comprehensive tool for microbial pangenome-wide association studies. Bioinformatics [Internet] 34:4310– 4312. Available from: 10.1093/bioinformatics/bty539

Lees JA, Harris SR, Tonkin-Hill G, Gladstone RA, Lo SW, Weiser JN, Corander J, Bentley SD, Croucher NJ. 2019. Fast and flexible bacterial genomic epidemiology with PopPUNK. Genome Res. [Internet] 29:304–316. Available from: 10.1101/gr.241455.118

Liu X, Xu Q, Yang X, Heng H, Yang C, Yang G, Peng M, Chan EW-C, Chen S. 2025. Capsular polysaccharide enables to evade phagocytosis by blocking host-bacteria interactions. mBio [Internet] 16:e0383824. Available from: 10.1128/mbio.03838-24

Løchen A, Truscott JE, Croucher NJ. 2022. Analysing pneumococcal invasiveness using Bayesian models of pathogen progression rates. PLoS Comput Biol [Internet] 18:e1009389. Available from: 10.1371/journal.pcbi.1009389

Lo SW, Mellor K, Cohen R, Alonso AR, Belman S, Kumar N, Hawkins PA, Gladstone RA, von Gottberg A, Veeraraghavan B, et al. 2022. Emergence of a multidrug-resistant and virulent Streptococcus pneumoniae lineage mediates serotype replacement after PCV13: an international whole-genome sequencing study. Lancet Microbe [Internet] 3:e735–e743. Available from: 10.1016/S2666-5247(22)00158-6

Lucidarme J, Lekshmi A, Parikh SR, Bray JE, Hill DM, Bratcher HB, Gray SJ, Carr AD, Jolley KA, Findlow J, et al. 2017. Frequent capsule switching in “ultra-virulent” meningococci - Are we ready for a serogroup B ST-11 complex outbreak? J. Infect. [Internet] 75:95–103. Available from: 10.1016/j.jinf.2017.05.015

Maiden MCJ, Jansen van Rensburg MJ, Bray JE, Earle SG, Ford SA, Jolley KA, McCarthy ND. 2013. MLST revisited: the gene-by-gene approach to bacterial genomics. Nat. Rev. Microbiol. [Internet] 11:728–736. Available from: 10.1038/nrmicro3093

Majewski J, Cohan FM. 1999. DNA sequence similarity requirements for interspecific recombination in Bacillus. Genetics [Internet] 153:1525–1533. Available from: 10.1093/genetics/153.4.1525

Majewski J, Zawadzki P, Pickerill P, Cohan FM, Dowson CG. 2000. Barriers to genetic exchange between bacterial species: Streptococcus pneumoniae transformation. J. Bacteriol. [Internet] 182:1016–1023. Available from: 10.1128/JB.182.4.1016-1023.2000

Marks LR, Reddinger RM, Hakansson AP. 2012. High levels of genetic recombination during nasopharyngeal carriage and biofilm formation in Streptococcus pneumoniae. mBio [Internet] 3. Available from: 10.1128/mBio.00200-12

Martins ER, Melo-Cristino J, Ramirez M. 2010. Evidence for Rare Capsular Switching in Streptococcus agalactiae. J. Bacteriol. [Internet]. Available from: https://journals.asm.org/doi/10.1128/jb.01130-09

Ma SX, Eichner H, Cammer M, Weiser JN. 2025. Inhibitory effect of capsule on natural transformation of. bioRxiv [Internet]. Available from: 10.1101/2025.05.07.652784

Mavroidi A, Aanensen DM, Godoy D, Skovsted IC, Kaltoft MS, Reeves PR, Bentley SD, Spratt BG. 2007. Genetic relatedness of the Streptococcus pneumoniae capsular biosynthetic loci. J. Bacteriol. [Internet] 189:7841–7855. Available from: 10.1128/JB.00836-07

Mazzamurro F, Chirakadavil JB, Durieux I, Poiré L, Plantade J, Ginevra C, Jarraud S, Wilharm G, Charpentier X, P C Rocha E. 2024. Intragenomic conflicts with plasmids and chromosomal mobile genetic elements drive the evolution of natural transformation within species. PLoS Biol [Internet] 22:e3002814. Available from: 10.1371/journal.pbio.3002814

Moore MR, Gertz RE Jr, Woodbury RL, Barkocy-Gallagher GA, Schaffner W, Lexau C, Gershman K, Reingold A, Farley M, Harrison LH, et al. 2008. Population snapshot of emergent Streptococcus pneumoniae serotype 19A in the United States, 2005. J. Infect. Dis. [Internet] 197:1016–1027. Available from: 10.1086/528996

Mostowy R, Croucher NJ, Hanage WP, Harris SR, Bentley S, Fraser C. 2014. Heterogeneity in the frequency and characteristics of homologous recombination in pneumococcal evolution. PLoS Genet. [Internet] 10:e1004300. Available from: 10.1371/journal.pgen.1004300

Mostowy RJ, Croucher NJ, De Maio N, Chewapreecha C, Salter SJ, Turner P, Aanensen DM, Bentley SD, Didelot X, Fraser C. 2017. Pneumococcal Capsule Synthesis Locus cps as Evolutionary Hotspot with Potential to Generate Novel Serotypes by Recombination. Mol. Biol. Evol. [Internet] 34:2537–2554. Available from: 10.1093/molbev/msx173

Mostowy RJ, Holt KE. 2018. Diversity-Generating Machines: Genetics of Bacterial Sugar-Coating. Trends Microbiol. [Internet] 26:1008–1021. Available from: 10.1016/j.tim.2018.06.006

Murray GGR, Hossain ASMM, Miller EL, Bruchmann S, Balmer AJ, Matuszewska M, Herbert J, Hadjirin NF, Mugabi R, Li G, et al. 2023. The emergence and diversification of a zoonotic pathogen from within the microbiota of intensively farmed pigs. Proc. Natl. Acad. Sci. U. S. A. [Internet] 120:e2307773120. Available from: 10.1073/pnas.2307773120

Nelson AL, Roche AM, Gould JM, Chim K, Ratner AJ, Weiser JN. 2007. Capsule enhances pneumococcal colonization by limiting mucus-mediated clearance. Infect Immun [Internet] 75:83–90. Available from: 10.1128/IAI.01475-06

Nguyen L-T, Schmidt HA, von Haeseler A, Minh BQ. 2015. IQ-TREE: a fast and effective stochastic algorithm for estimating maximum-likelihood phylogenies. Mol. Biol. Evol. [Internet] 32:268–274. Available from: 10.1093/molbev/msu300

Ondov BD, Treangen TJ, Melsted P, Mallonee AB, Bergman NH, Koren S, Phillippy AM. 2016. Mash: fast genome and metagenome distance estimation using MinHash. Genome Biol. [Internet] 17:132. Available from: 10.1186/s13059-016-0997-x

Pai R, Gertz RE, Beall B. 2006. Sequential multiplex PCR approach for determining capsular serotypes of Streptococcus pneumoniae isolates. J. Clin. Microbiol. [Internet] 44:124–131. Available from: 10.1128/JCM.44.1.124-131.2006

Paradis E, Claude J, Strimmer K. 2004. APE: Analyses of Phylogenetics and Evolution in R language. Bioinformatics [Internet] 20:289–290. Available from: 10.1093/bioinformatics/btg412

Paton JC, Trappetti C. 2019. *Streptococcus pneumoniae* capsular polysaccharide. Microbiol. Spectr. [Internet] 7. Available from: https://journals.asm.org/doi/10.1128/microbiolspec.GPP3-0019-2018

Pearce BJ, Iannelli F, Pozzi G. 2002. Construction of new unencapsulated (rough) strains of Streptococcus pneumoniae. Res Microbiol [Internet] 153:243–247. Available from: 10.1016/s0923-2508(02)01312-8

Pillai DR, Shahinas D, Buzina A, Pollock RA, Lau R, Khairnar K, Wong A, Farrell DJ, Green K, McGeer A, et al. 2009. Genome-wide dissection of globally emergent multi-drug resistant serotype 19A Streptococcus pneumoniae. BMC Genomics [Internet] 10:642. Available from: 10.1186/1471-2164-10-642

Popa O, Dagan T. 2011. Trends and barriers to lateral gene transfer in prokaryotes. Curr. Opin. Microbiol. [Internet] 14:615–623. Available from: 10.1016/j.mib.2011.07.027

Popa O, Hazkani-Covo E, Landan G, Martin W, Dagan T. 2011. Directed networks reveal genomic barriers and DNA repair bypasses to lateral gene transfer among prokaryotes. Genome Res. [Internet] 21:599–609. Available from: 10.1101/gr.115592.110

Power RA, Parkhill J, de Oliveira T. 2017. Microbial genome-wide association studies: lessons from human GWAS. Nat. Rev. Genet. [Internet] 18:41–50. Available from: 10.1038/nrg.2016.132

Rappuoli R, De Gregorio E, Costantino P. 2019. On the mechanisms of conjugate vaccines. Proc. Natl. Acad. Sci. U. S. A. [Internet] 116:14–16. Available from: 10.1073/pnas.1819612116

Revell LJ. 2012. phytools: an R package for phylogenetic comparative biology (and other things). Methods in Ecology and Evolution [Internet] 3:217–223. Available from: 10.1111/j.2041-210x.2011.00169.x

Riley MA, Gordon DM. 1999. The ecological role of bacteriocins in bacterial competition. Trends Microbiol [Internet] 7:129–133. Available from: 10.1016/s0966-842x(99)01459-6

Ritchie ND, Mitchell TJ, Evans TJ. 2012. What is different about serotype 1 pneumococci? Future Microbiol. [Internet] 7:33–46. Available from: 10.2217/fmb.11.146

Sheppard CL, Groves N, Andrews N, Litt DJ, Fry NK, Southern J, Miller E. 2019. The Genomics of Carriage Isolates from UK Children and Their Household Contacts, Pre-PCV7 to Post-PCV13. Genes [Internet] 10. Available from: 10.3390/genes10090687

Song J-H, Ko KS, Lee J-Y, Baek JY, Oh WS, Yoon HS, Jeong J-Y, Chun J. 2005. Identification of essential genes in Streptococcus pneumoniae by allelic replacement mutagenesis. Mol Cells [Internet] 19:365–374. Available from: https://www.ncbi.nlm.nih.gov/pubmed/15995353

Soundararajan M, von Bünau R, Oelschlaeger TA. 2019. K5 Capsule and Lipopolysaccharide Are Important in Resistance to T4 Phage Attack in Probiotic Strain Nissle 1917. Front. Microbiol. [Internet] 10:2783. Available from: 10.3389/fmicb.2019.02783

Tettelin H, Nelson KE, Paulsen IT, Eisen JA, Read TD, Peterson S, Heidelberg J, DeBoy RT, Haft DH, Dodson RJ, et al. 2001. Complete genome sequence of a virulent isolate of Streptococcus pneumoniae. Science [Internet] 293:498–506. Available from: 10.1126/science.1061217

Trzciński K, Li Y, Weinberger DM, Thompson CM, Cordy D, Bessolo A, Malley R, Lipsitch M. 2015. Effect of Serotype on Pneumococcal Competition in a Mouse Colonization Model. mBio [Internet] 6:e00902–e00915. Available from: 10.1128/mBio.00902-15

Turner SD. qqman: an R package for visualizing GWAS results using Q-Q and manhattan plots. Available from: 10.1101/005165

Tyler S, Tsang R. 2004. Genetic analysis of Canadian isolates of C:2a:P1.2,5 and B:2a:P1.2,5 Neisseria meningitidis strains belonging to the hypervirulent clone of ET-15. Can. J. Microbiol. [Internet] 50:433–443. Available from: 10.1139/w04-024

Wahl B, O’Brien KL, Greenbaum A, Majumder A, Liu L, Chu Y, Lukšić I, Nair H, McAllister DA, Campbell H, et al. 2018. Burden of Streptococcus pneumoniae and Haemophilus influenzae type b disease in children in the era of conjugate vaccines: global, regional, and national estimates for 2000-15. Lancet Glob Health [Internet] 6:e744–e757. Available from: 10.1016/S2214-109X(18)30247-X

Wang J, Feng Y, Zong Z. 2023. The Origins of ST11 KL64 Klebsiella pneumoniae: a Genome-Based Study. Microbiol Spectr [Internet] 11:e0416522. Available from: 10.1128/spectrum.04165-22

Weinberger DM, Trzciński K, Lu Y-J, Bogaert D, Brandes A, Galagan J, Anderson PW, Malley R, Lipsitch M. 2009. Pneumococcal capsular polysaccharide structure predicts serotype prevalence. PLoS Pathog. [Internet] 5:e1000476. Available from: 10.1371/journal.ppat.1000476

Weiser JN, Kapoor M. 1999. Effect of intrastrain variation in the amount of capsular polysaccharide on genetic transformation of Streptococcus pneumoniae: implications for virulence studies of encapsulated strains. Infect Immun [Internet] 67:3690–3692. Available from: 10.1128/IAI.67.7.3690-3692.1999

Whitfield C, Wear SS, Sande C. 2020. Assembly of Bacterial Capsular Polysaccharides and Exopolysaccharides. Annu. Rev. Microbiol. [Internet] 74:521–543. Available from: 10.1146/annurev-micro-011420-075607

Wood DE, Salzberg SL. 2014. Kraken: ultrafast metagenomic sequence classification using exact alignments. Genome Biol. [Internet] 15:R46. Available from: 10.1186/gb-2014-15-3-r46

Wyres KL, Gorrie C, Edwards DJ, Wertheim HFL, Hsu LY, Van Kinh N, Zadoks R, Baker S, Holt KE. 2015. Extensive Capsule Locus Variation and Large-Scale Genomic Recombination within the Klebsiella pneumoniae Clonal Group 258. Genome Biol. Evol. [Internet] 7:1267–1279. Available from: 10.1093/gbe/evv062

Wyres KL, Lambertsen LM, Croucher NJ, McGee L, von Gottberg A, Liñares J, Jacobs MR, Kristinsson KG, Beall BW, Klugman KP, et al. 2013. Pneumococcal capsular switching: a historical perspective. J. Infect. Dis. [Internet] 207:439–449. Available from: 10.1093/infdis/jis703

Yother J, McDaniel LS, Briles DE. 1986. Transformation of encapsulated Streptococcus pneumoniae. J Bacteriol [Internet] 168:1463–1465. Available from: 10.1128/jb.168.3.1463-1465.1986

