## Supplementary Information for "The global evolutionary dynamics of natural capsule switching in a human-restricted bacterial pathogen"

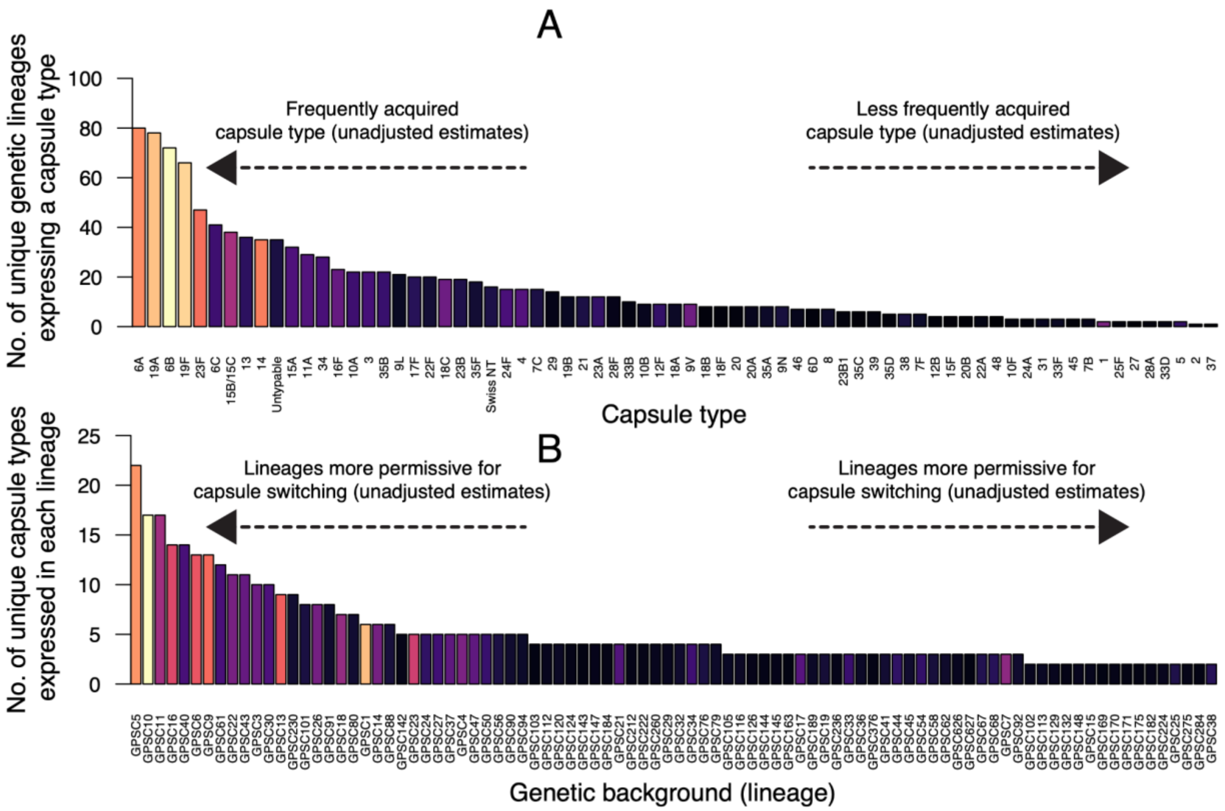


**Supplementary Fig. 1. Genetic and non-genetic factors are associated with capsule switching in the global phylogeny**. **(A)** The bar chart shows the unnormalized number of distinct genetic backgrounds that express a particular capsule type. The specific counts shown in the bar chart are provided in **Supplementary Data 2**. **(B)** Bar chart showing the unnormalized number of unique capsule types expressed in a single genetic background or lineage. For clarity, only the top 50 genetic backgrounds or GPSCs expressing the most capsule types are shown in the bar chart. The complete list of genetic backgrounds and the number of unique capsule types they contain is available in **Supplementary Data 4**. Panels D and E are based on only capsule types and lineages with at least ten sequenced genomes in the dataset, and the color of the bars in the bar plots corresponds to the number of genomes included in the analysis for the capsule type or lineage – the lighter the color, the more the genomes **(Supplementary Data 1)**.

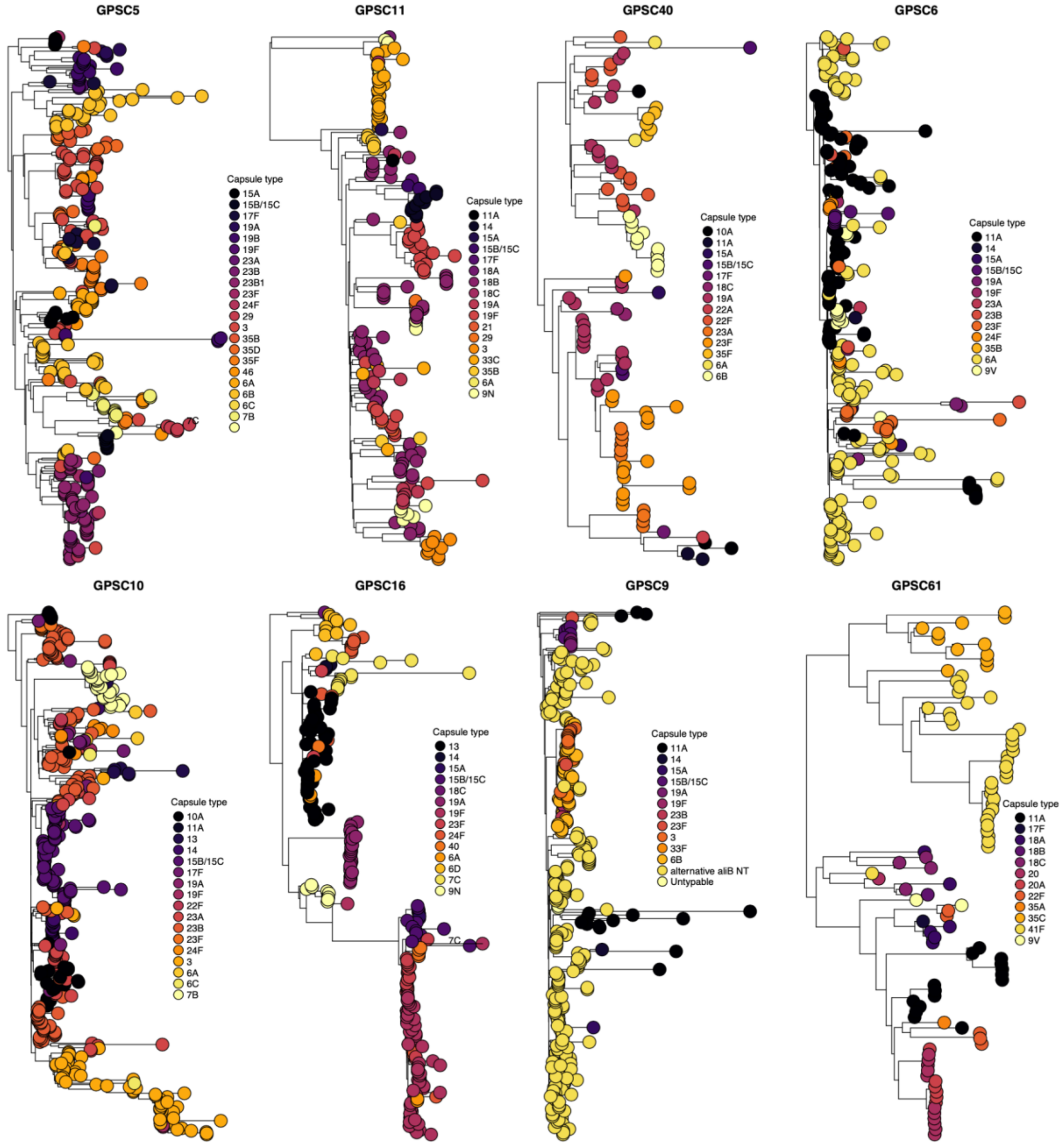


**Supplementary Fig. 2. Whole-genome phylogenies of pneumococcal lineages expressing many capsule types**. A maximum likelihood phylogeny showing examples of GPSC lineages expressing many capsule types, indicates a genetic background potentially more permissive to capsule switching.

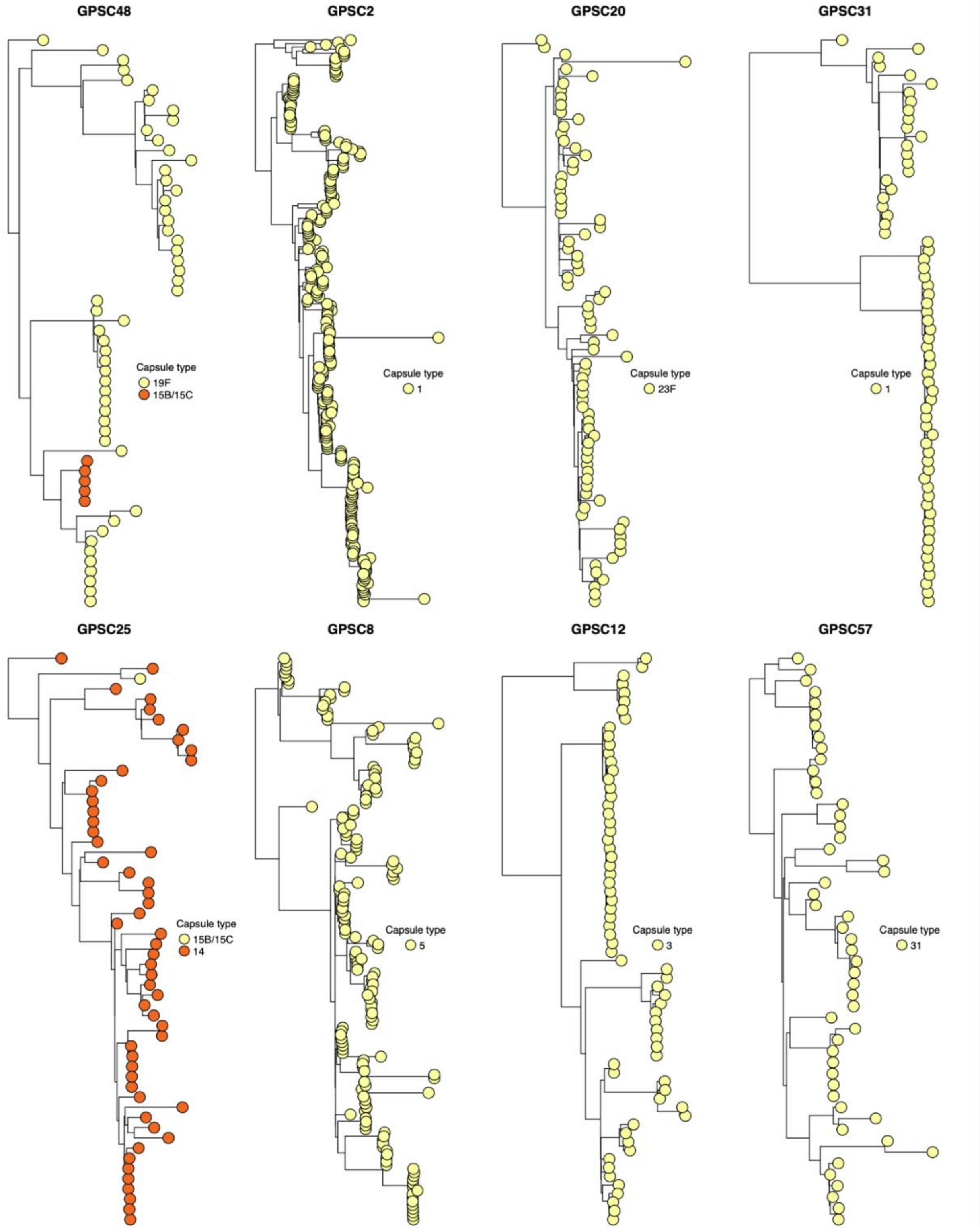


**Supplementary Fig. 3. Whole-genome phylogenies of pneumococcal lineages expressing few capsule types**. A maximum likelihood phylogeny showing examples of GPSC lineages expressing many capsule types, indicates a genetic background potentially more permissive to capsule switching.

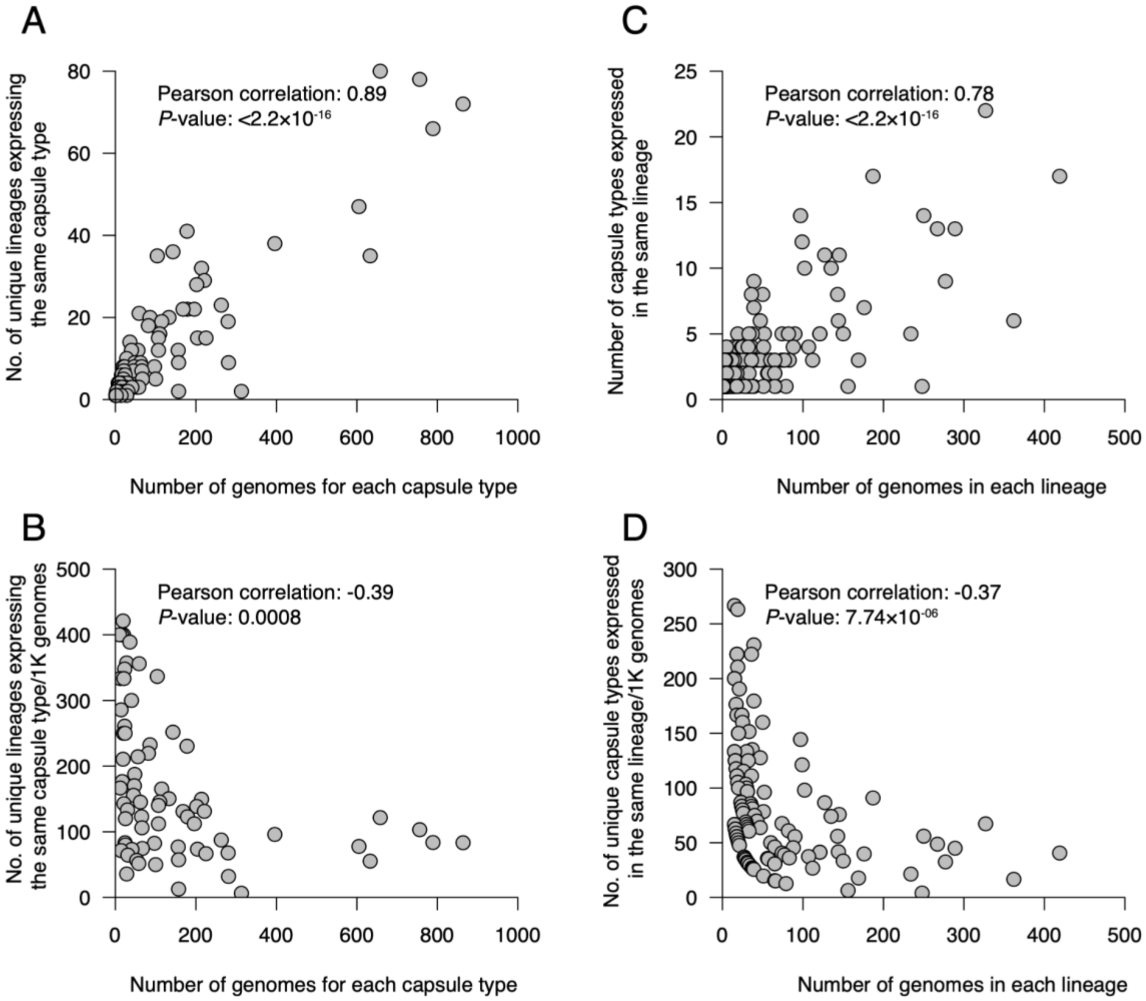


**Supplementary Fig. 4. Normalized and unnormalized counts of the number of lineages expressing each capsule type and number of pneumococcal capsule types expressed in a single lineage**. **(A)** A scatter plot shows the association between the number of lineages expressing each capsule type (i.e., capsule acquisitions) and the number of genomes of each capsule type. The strong correlation between these estimates suggested the need to normalize the estimates by the number of genomes as oversampled capsule types show expression across more lineages than those undersampled. **(B)** A scatter plot showing normalized estimates of the number of lineages expressing each capsule type per 1,000 genomes compared to the number of genomes associated with each capsule type. Only capsule types with at least 10 genomes in the dataset were normalized. **(C)** A scatter plot shows the association between the number of capsule types expressed in each lineage and the number of genomes corresponding to each lineage. The strong correlation between these estimates suggested the need to normalize the estimates by the number of genomes as oversampled capsule types show expression across more lineages than those undersampled. **(D)** A scatter plot showing normalized estimates of the number of capsule types expressed in each lineage per 1,000 genomes compared to the number of genomes in each lineage. Only lineages with at least 10 genomes in the dataset were normalized.

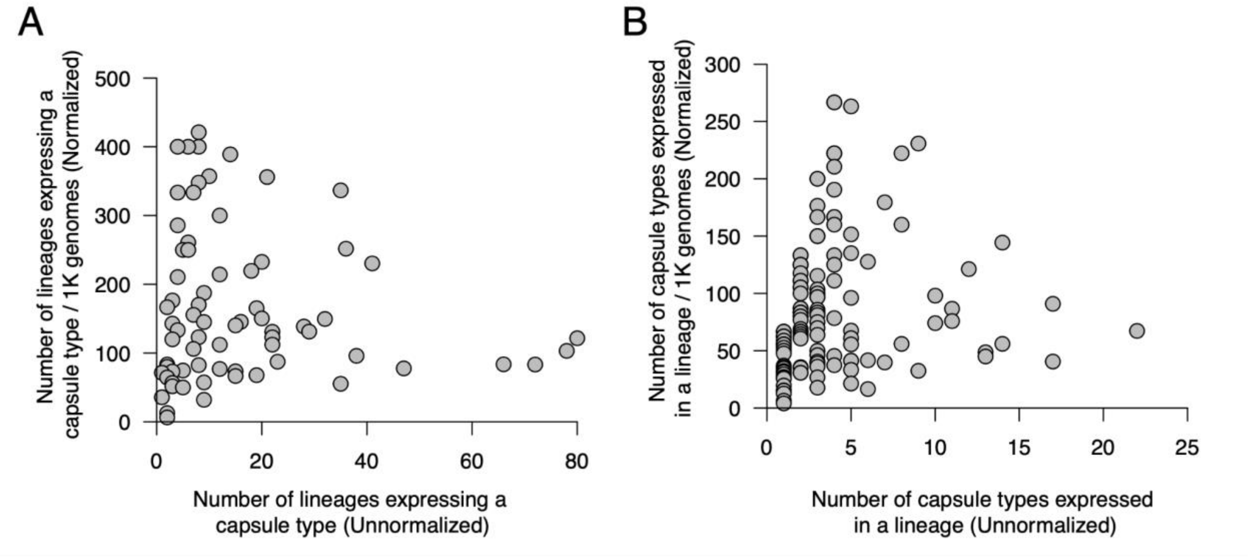


**Supplementary Fig. 5. Comparison of the normalized and unnormalized capsule switching estimates for each capsule type and genetic backgrounds**. **(A)** A scatter plot showing the normalized number of genetic backgrounds or lineages expressing each capsule type per 1,000 genomes and the unnormalized number of lineages expressing each capsule type. **(B)** A scatter plot showing the normalized number of capsule types expressed in each genetic background or lineage per 1,000 genomes and the unnormalized number of capsule types expressed in each lineage.

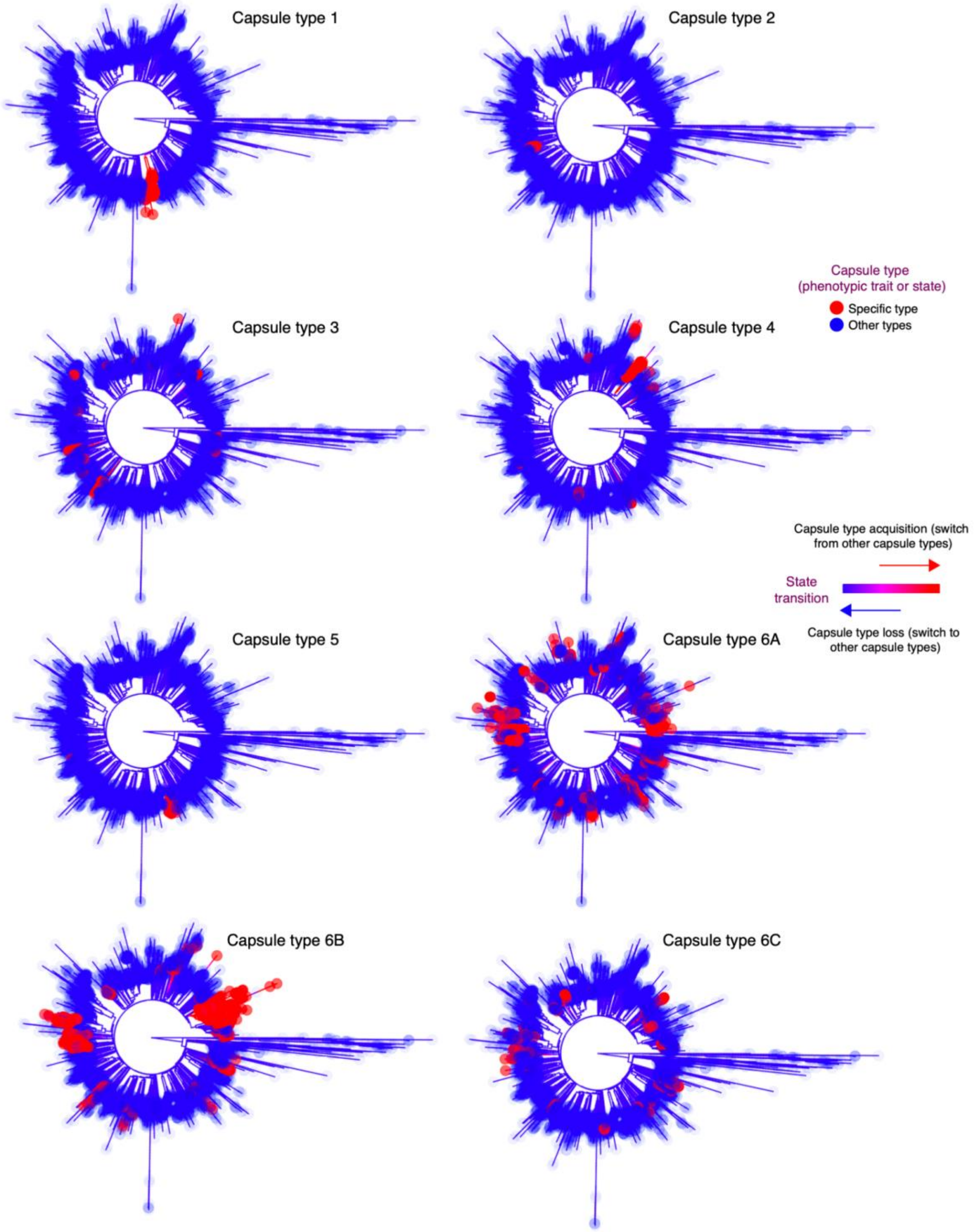


**Supplementary Fig. 6. Mapping the evolution of individual capsule types across the whole-genome-based pneumococcal phylogeny using ancestral state reconstruction.** Rooted global whole-genome maximum-likelihood phylogenies showing reconstructed capsule acquisition and loss patterns.

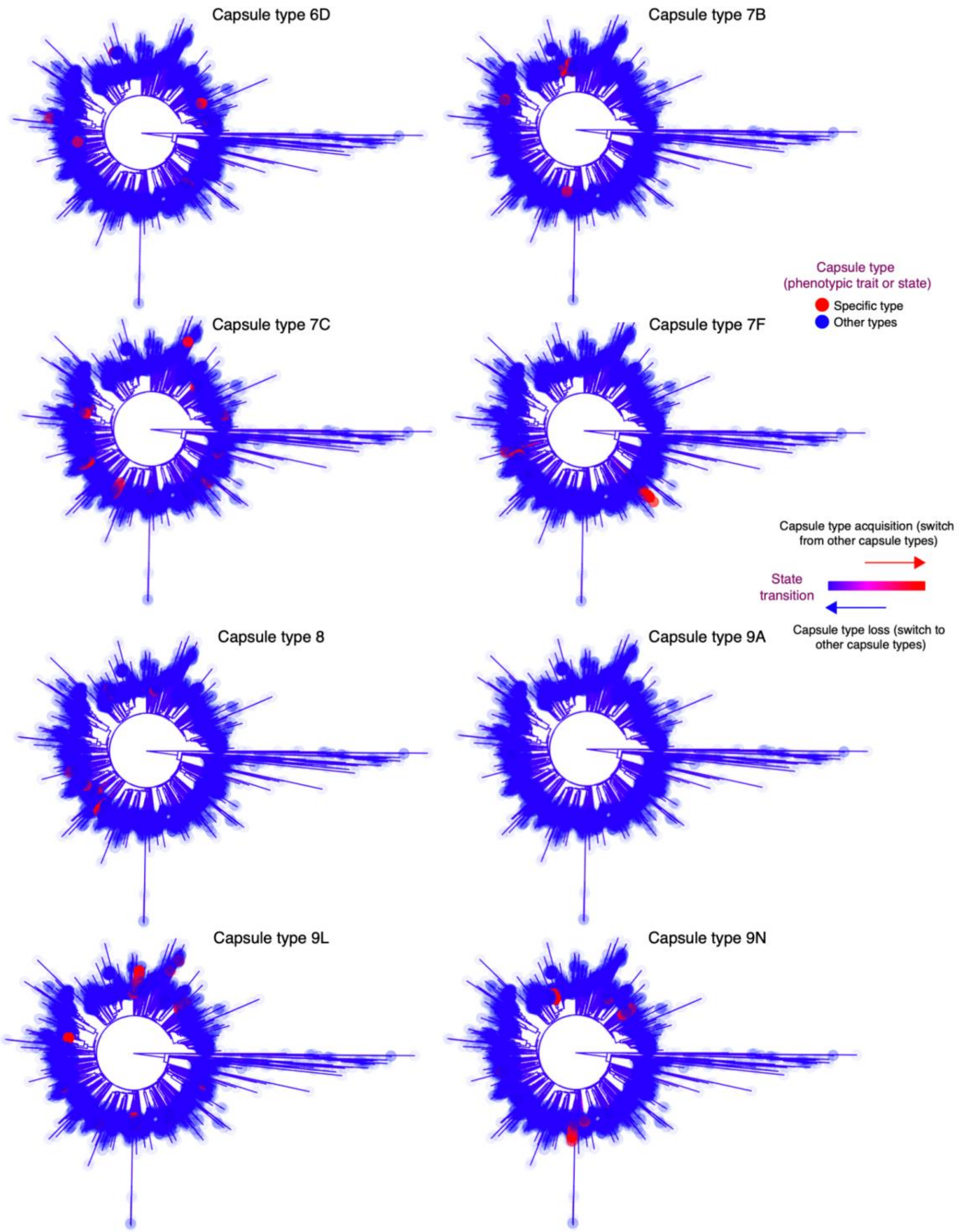


**Supplementary Fig. 6 (continued). Mapping the evolution of individual capsule types across the whole-genome-based pneumococcal phylogeny using ancestral state reconstruction.** Rooted global whole-genome maximum-likelihood phylogenies showing reconstructed capsule acquisition and loss patterns.

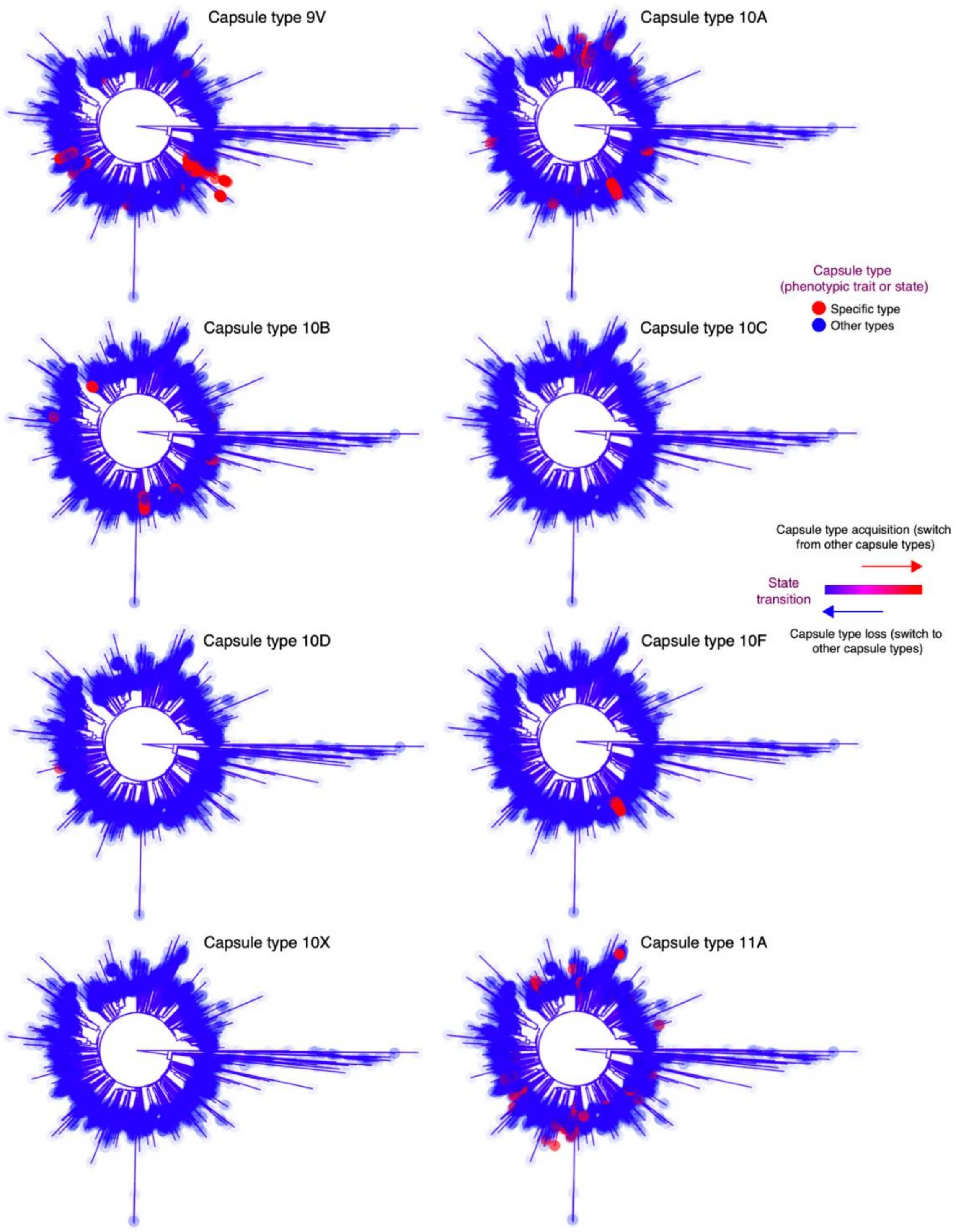


**Supplementary Fig. 6 (continued). Mapping the evolution of individual capsule types across the whole-genome-based pneumococcal phylogeny using ancestral state reconstruction.** Rooted global whole-genome maximum-likelihood phylogenies showing reconstructed capsule acquisition and loss patterns.

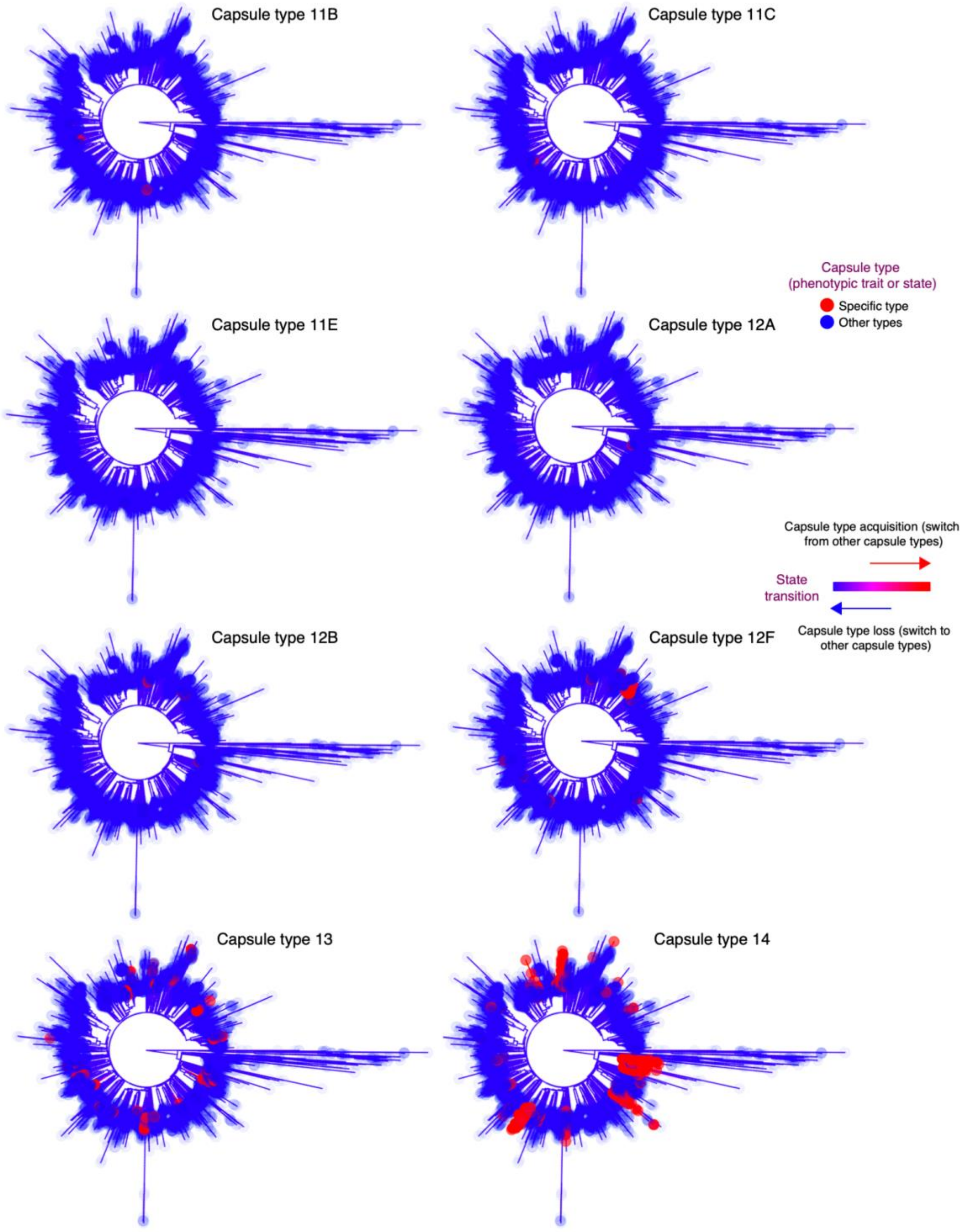


**Supplementary Fig. 6 (continued). Mapping the evolution of individual capsule types across the whole-genome-based pneumococcal phylogeny using ancestral state reconstruction.** Rooted global whole-genome maximum-likelihood phylogenies showing reconstructed capsule acquisition and loss patterns.

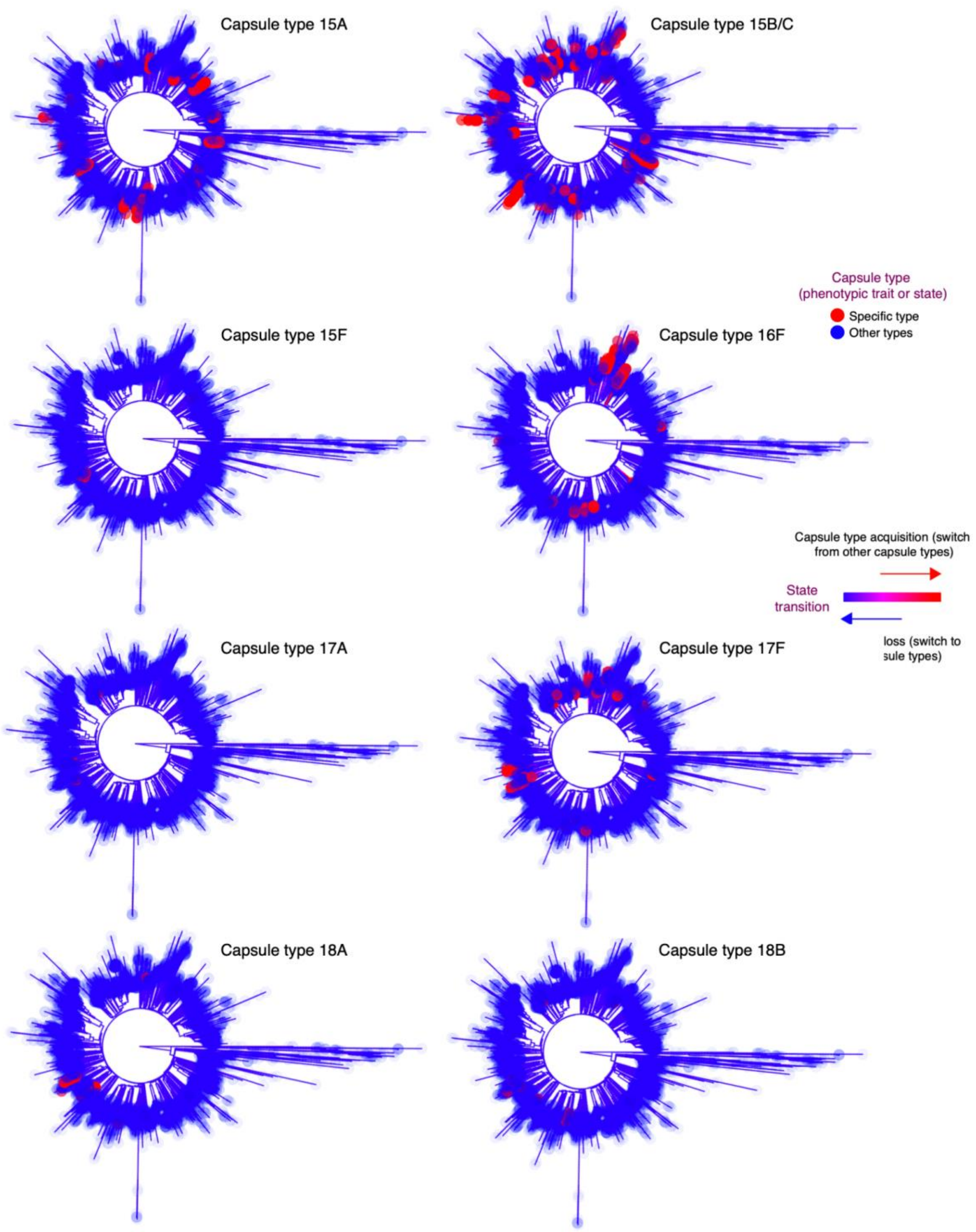


**Supplementary Fig. 6 (continued). Mapping the evolution of individual capsule types across the whole-genome-based pneumococcal phylogeny using ancestral state reconstruction.** Rooted global whole-genome maximum-likelihood phylogenies showing reconstructed capsule acquisition and loss patterns.

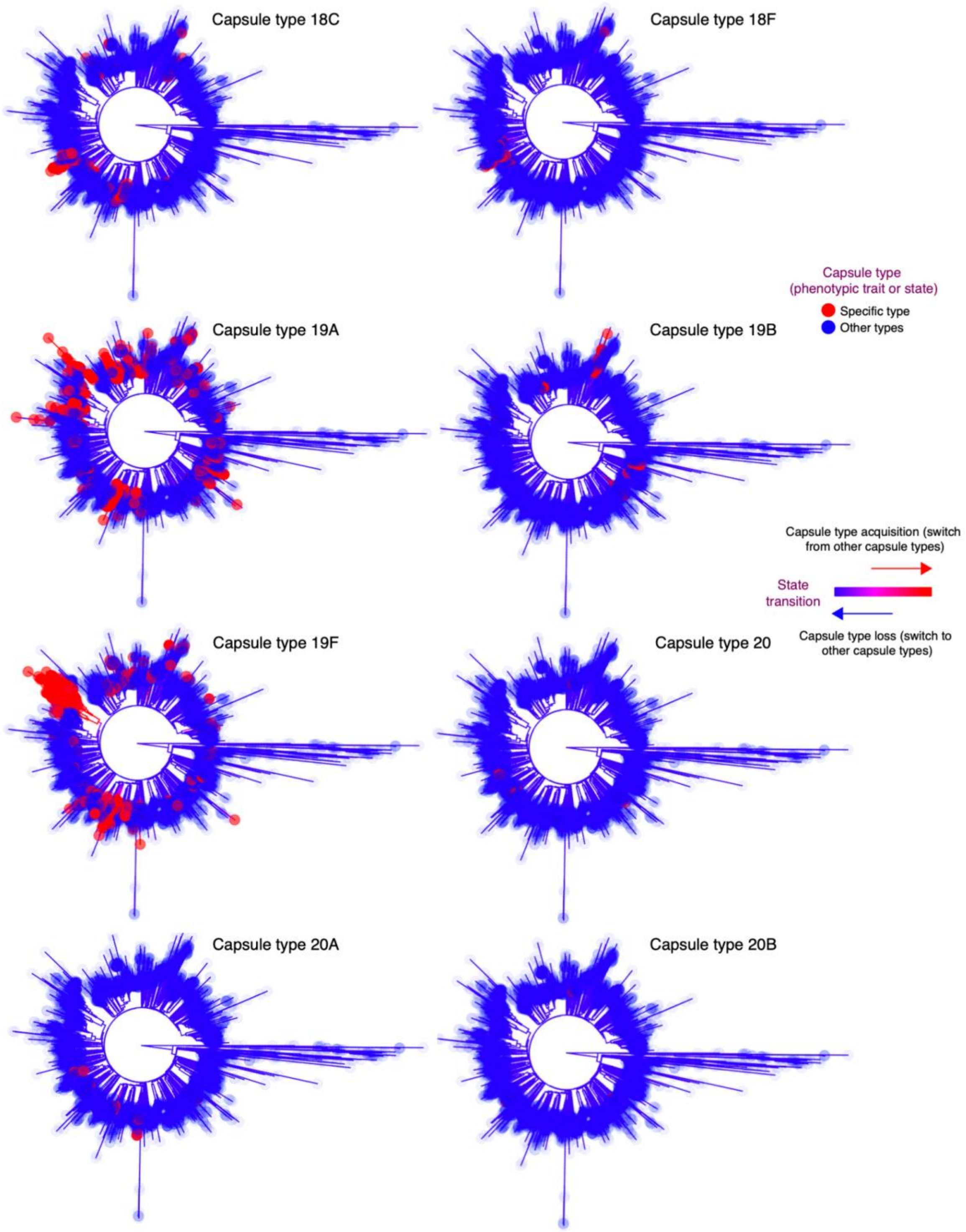


**Supplementary Fig. 6 (continued). Mapping the evolution of individual capsule types across the whole-genome-based pneumococcal phylogeny using ancestral state reconstruction.** Rooted global whole-genome maximum-likelihood phylogenies showing reconstructed capsule acquisition and loss patterns.

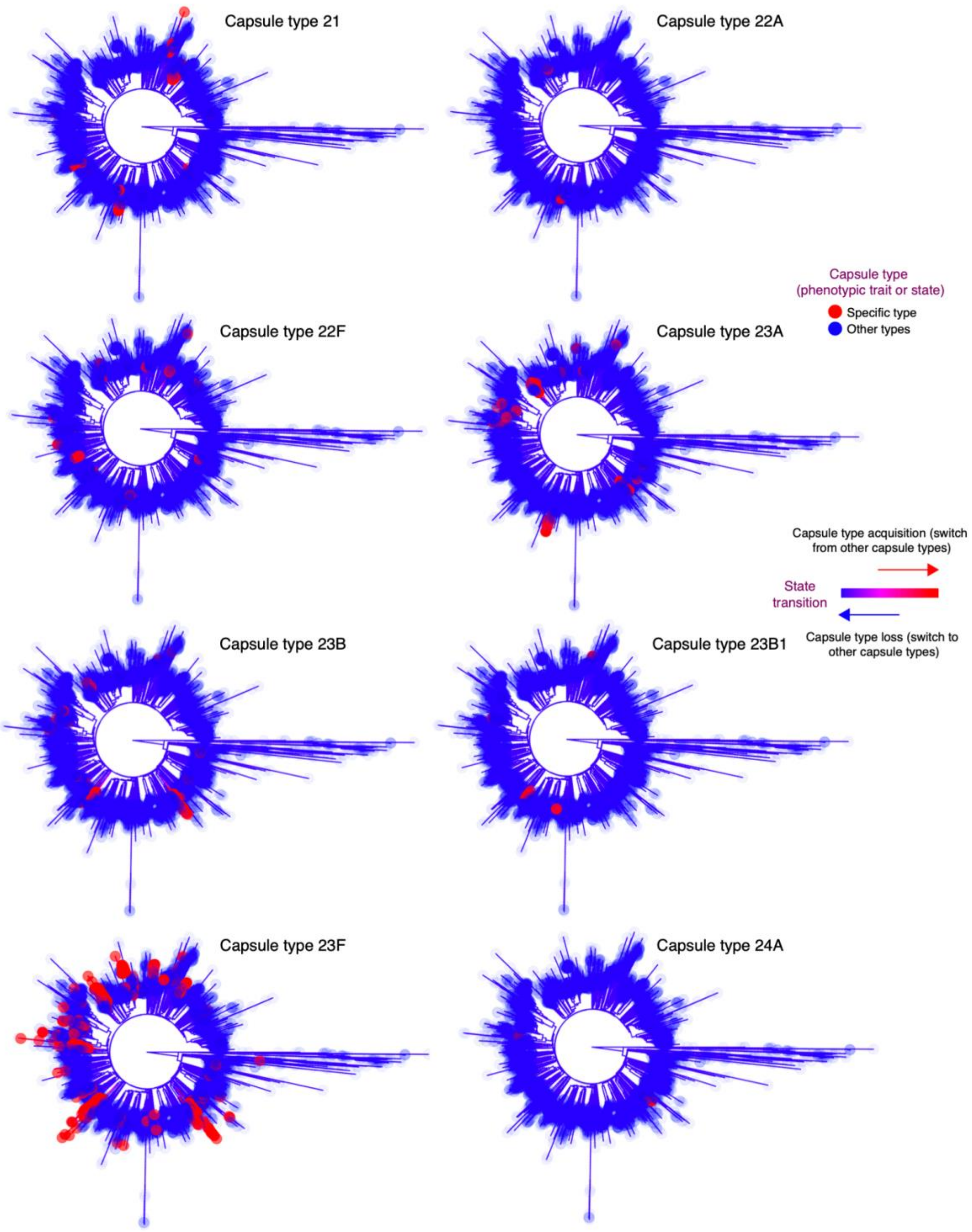


**Supplementary Fig. 6 (continued). Mapping the evolution of individual capsule types across the whole-genome-based pneumococcal phylogeny using ancestral state reconstruction.** Rooted global whole-genome maximum-likelihood phylogenies showing reconstructed capsule acquisition and loss patterns.

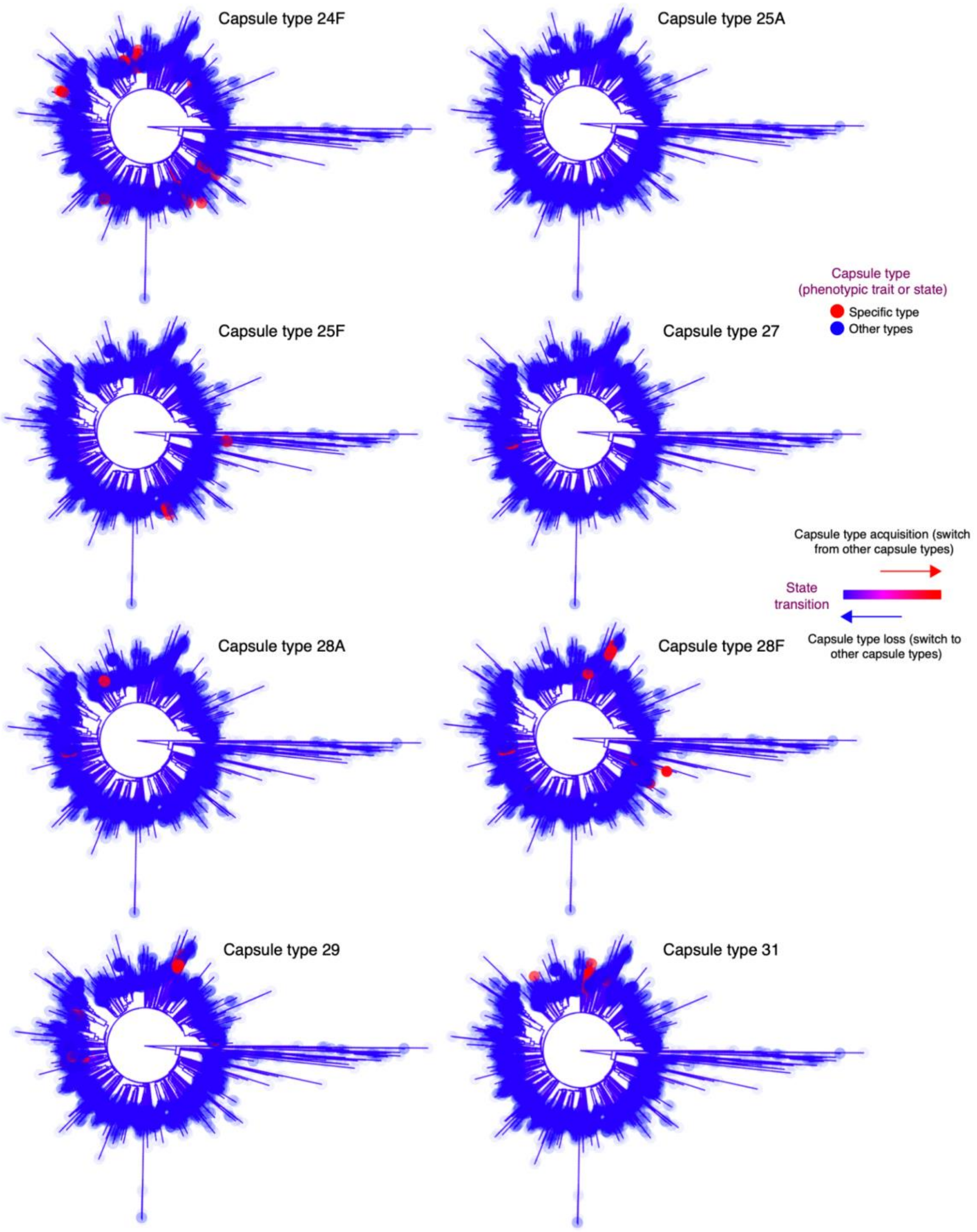


**Supplementary Fig. 6 (continued). Mapping the evolution of individual capsule types across the whole-genome-based pneumococcal phylogeny using ancestral state reconstruction.** Rooted global whole-genome maximum-likelihood phylogenies showing reconstructed capsule acquisition and loss patterns.

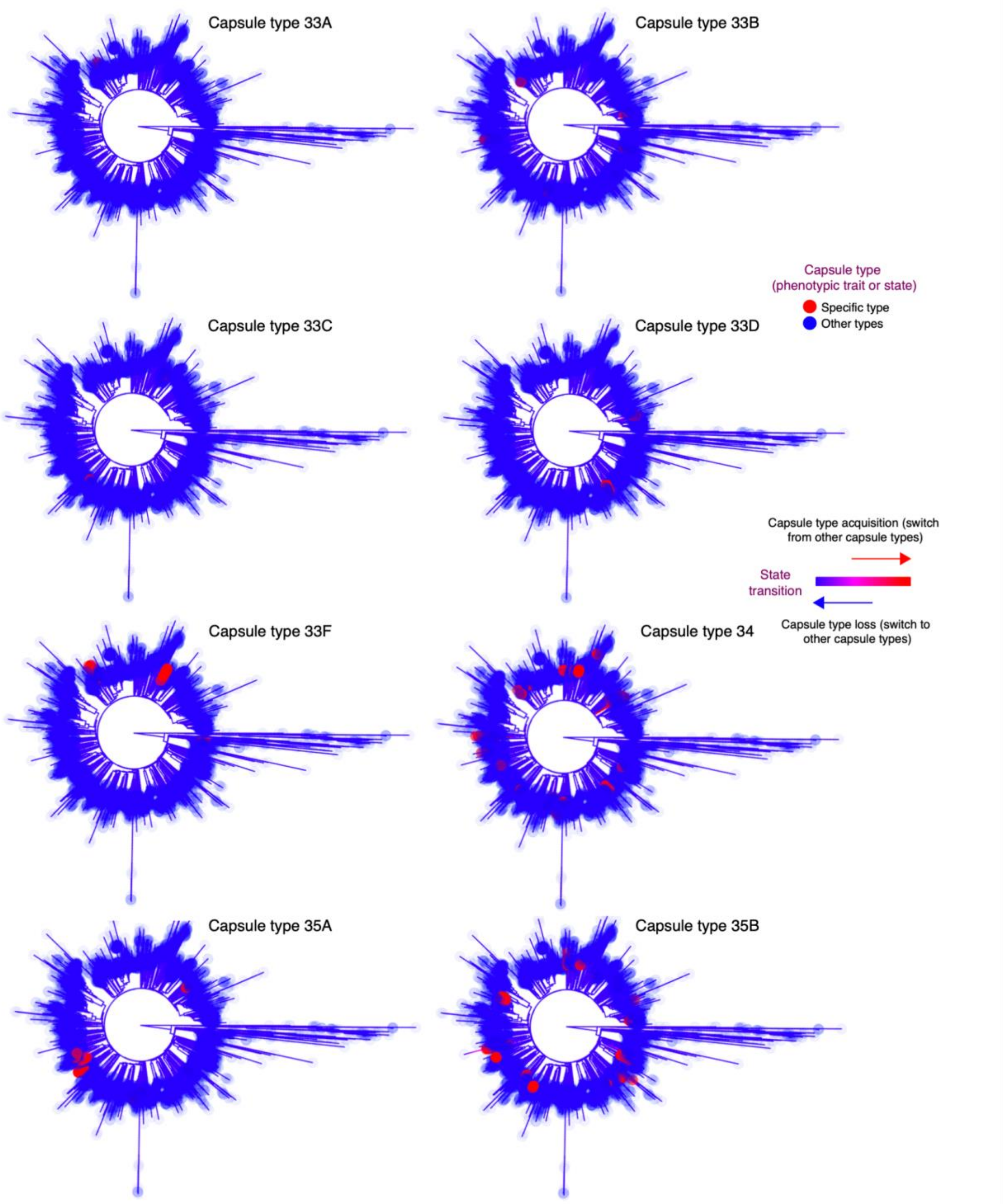


**Supplementary Fig. 6 (continued). Mapping the evolution of individual capsule types across the whole-genome-based pneumococcal phylogeny using ancestral state reconstruction.** Rooted global whole-genome maximum-likelihood phylogenies showing reconstructed capsule acquisition and loss patterns.

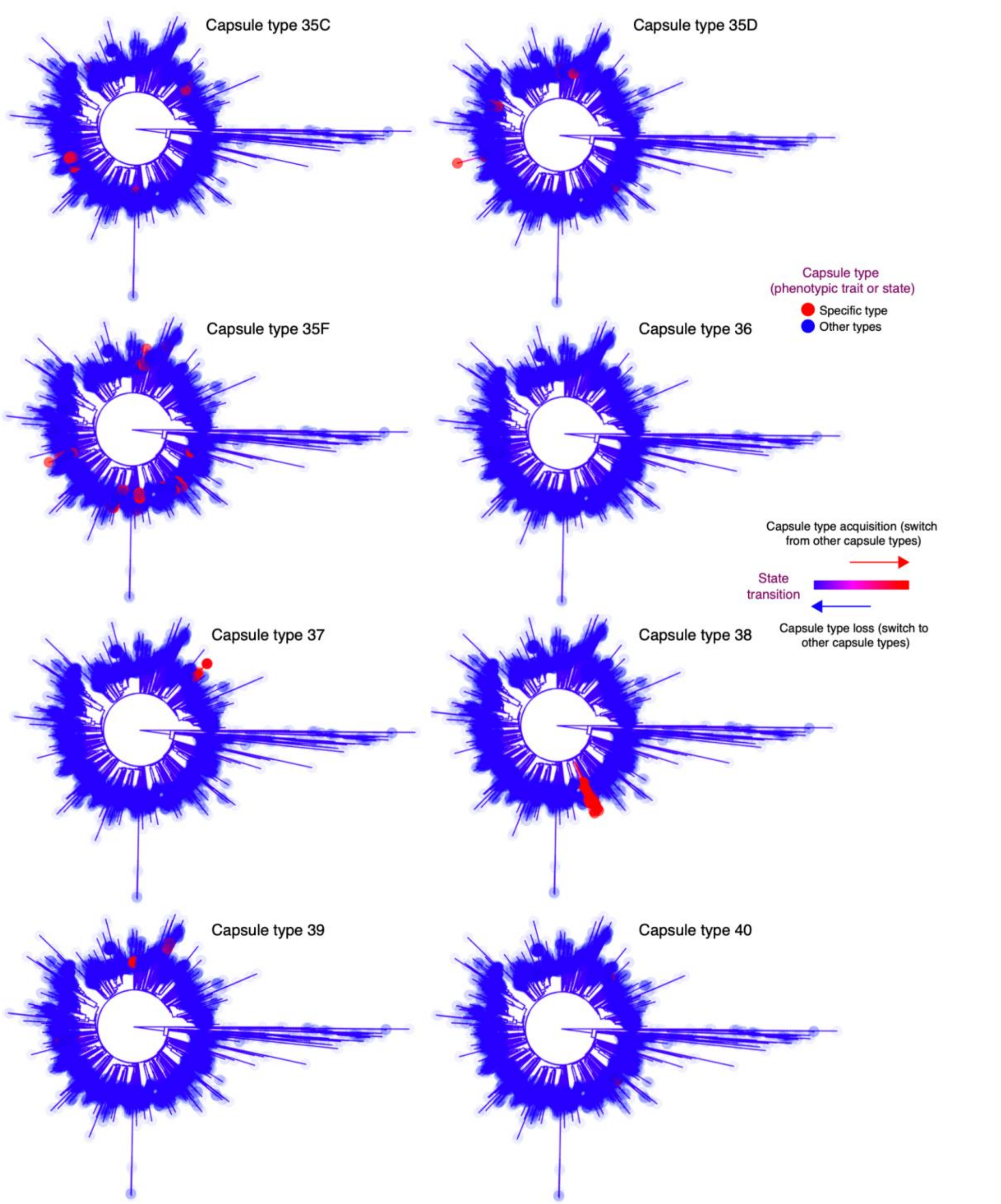


**Supplementary Fig. 6 (continued). Mapping the evolution of individual capsule types across the whole-genome-based pneumococcal phylogeny using ancestral state reconstruction.** Rooted global whole-genome maximum-likelihood phylogenies showing reconstructed capsule acquisition and loss patterns.

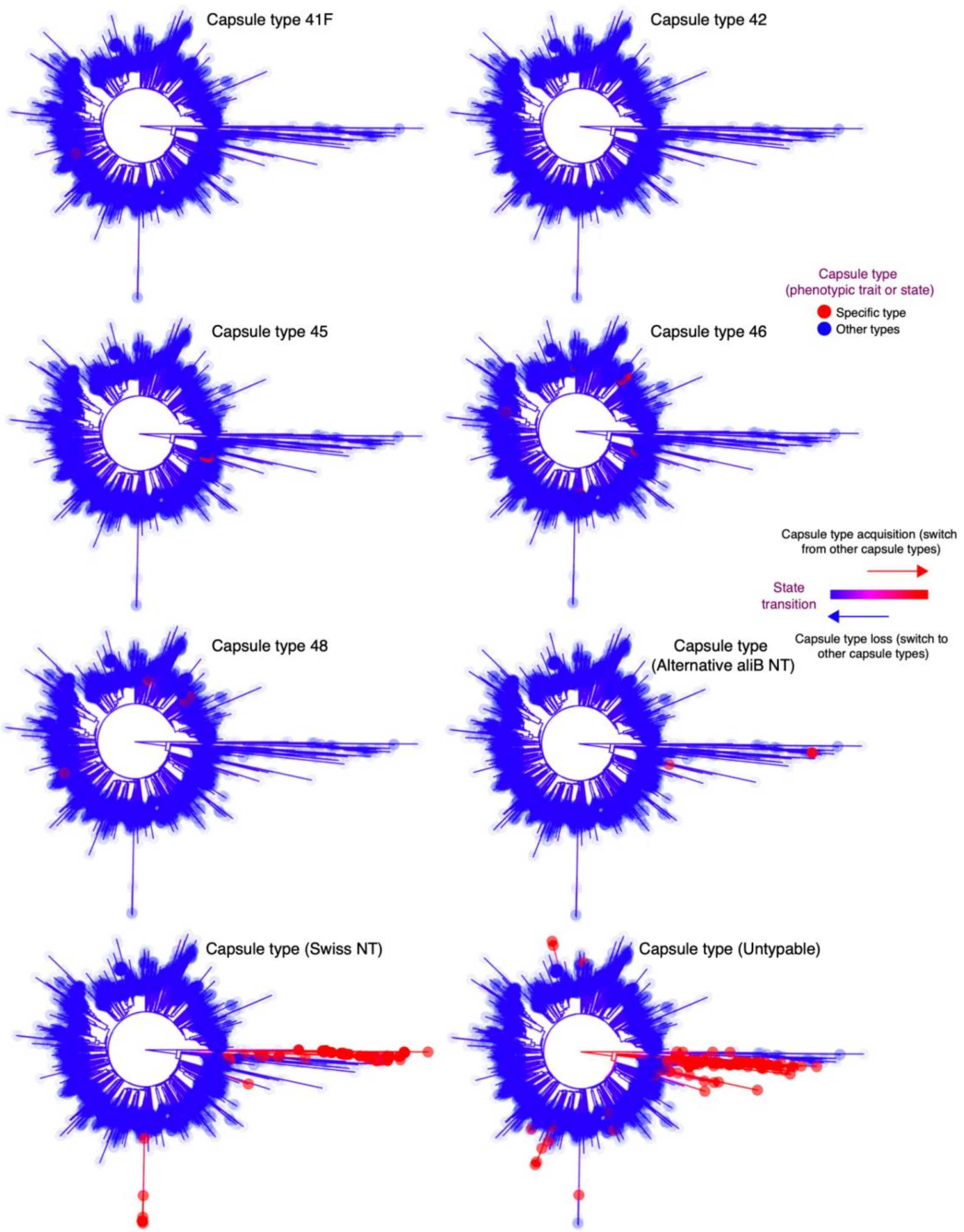


**Supplementary Fig. 6 (continued). Mapping the evolution of individual capsule types across the whole-genome-based pneumococcal phylogeny using ancestral state reconstruction.** Rooted global whole-genome maximum-likelihood phylogenies showing reconstructed capsule acquisition and loss patterns.

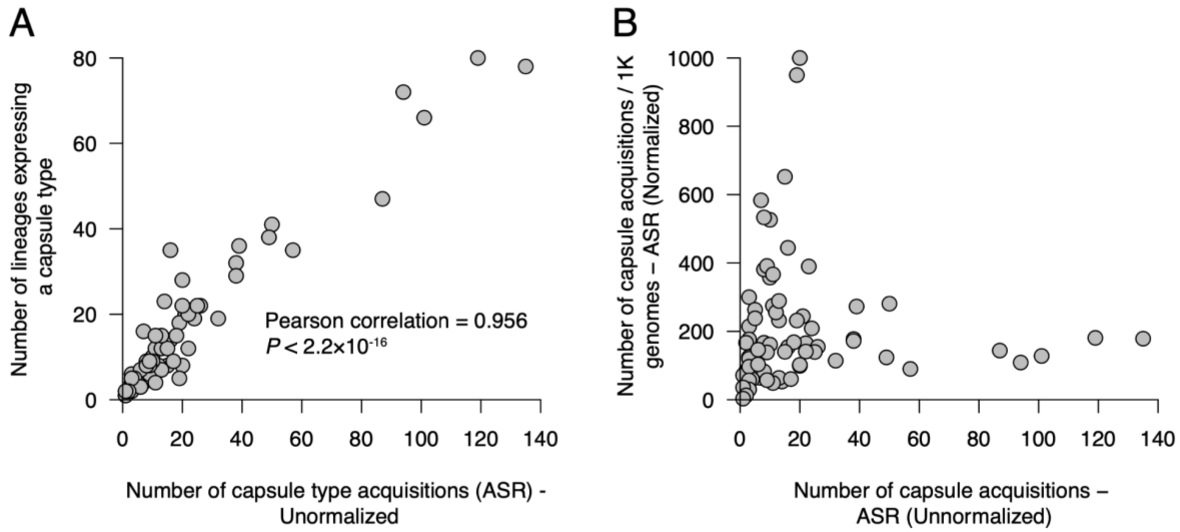


**Supplementary Fig. 7. Estimates of pneumococcal capsule acquisition**. **(A)** A scatter plot showing the number of acquisitions of capsule types based on counting the number of lineages expressing each capsule type and the number of capsule type acquisitions based on the ancestral state reconstruction (ASR) approach. **(B)** A scatter plot comparing the unnormalized and normalized number of capsule acquisitions in the phylogeny using the ASR approach.

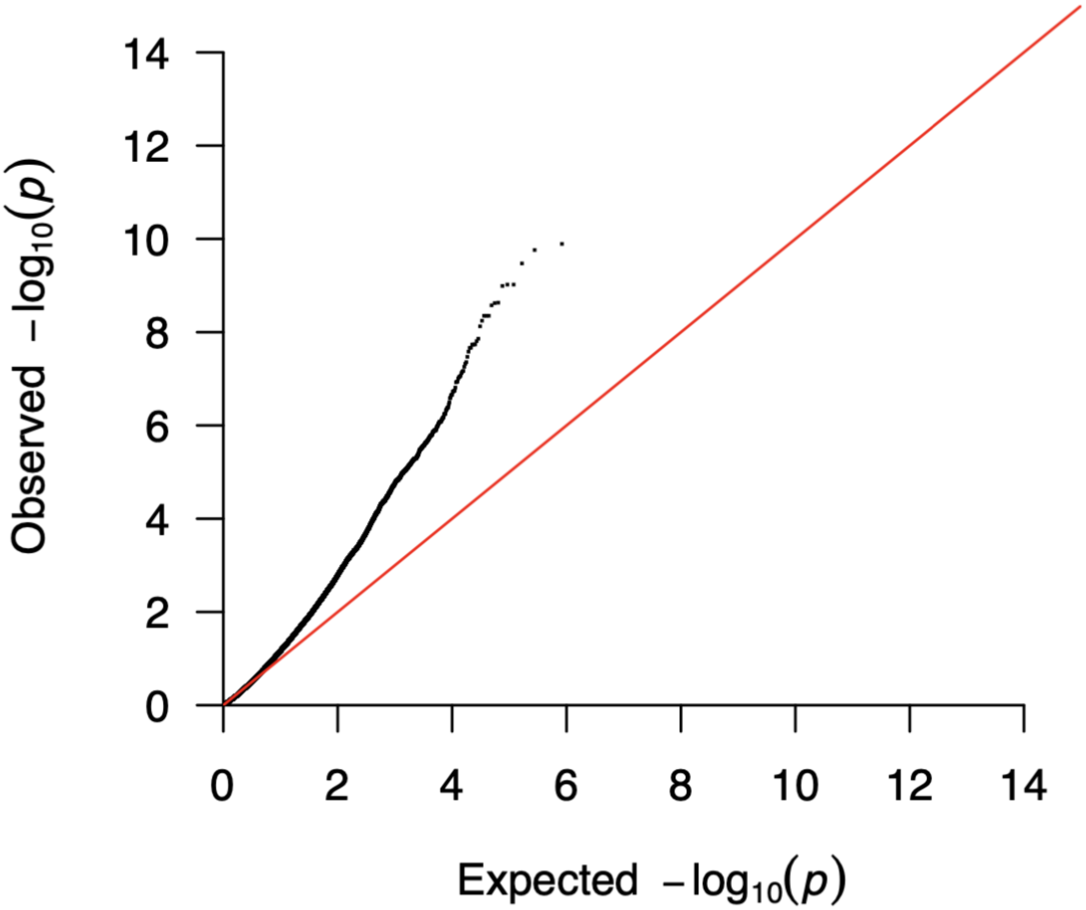


**Supplementary Fig. 8. Quantile-quantile (QQ) plot for the GWAS analysis to identify genomic loci associated with the propensity of pneumococcal lineages to capsule switching**. The QQ plot shows the relationship between the observed and expected statistical significance of the GWAS hits.
